# H2AK119ub dynamics controls hair follicle stem cell quiescence

**DOI:** 10.1101/2024.10.10.617646

**Authors:** Pooja Flora, Meng Yen Li, Yudong Zhou, Maria Mercédes, Xiang Yu Zheng, Phillip M. Galbo, Deyou Zheng, Elena Ezhkova

## Abstract

The transition of stem cells from a quiescent state to an active state is a finely tuned process that requires the dismantling of the quiescence program and the establishment of a cell cycle-promoting transcriptional landscape. Whether epigenetic processes control stem cell states to promote the regeneration of adult tissues remains elusive. In this study, we show that a repressive histone modification, H2AK119ub, is dynamic between quiescent and active hair follicle stem cells (HFSCs) in the adult murine skin. Ablation of H2AK119ub in HFSCs leads to impaired quiescence leading to premature activation and an eventual exhaustion of HFSC pool. Transcriptional and chromatin studies revealed that H2AK119ub directly represses a proliferation promoting transcriptional program in the HFSCs to preserve quiescence. Lastly, we identify that the inhibitory FGF signaling produced by the hair follicle niche keratinocytes maintains H2AK119ub in quiescent HFSCs. Together, these findings reveal that a repressive histone mark, H2AK119ub, is under the dynamic regulation of inhibitory niche signaling to prevent the untimely establishment of an activated state to preserve SC function and longevity.

## Main

Adult resident stem cells (SCs) are defined by their long-lived status, capacity to give rise to differentiating daughter cells and their ability to retain their identity by undergoing self-renewal^1,2^. While in some tissues, SCs engage in repetitive regeneration events to allow for rapid tissue turnover, in other tissues, they remain in a prolonged state of quiescence^3,4^. Dysregulation of this balance between SC quiescence and activation leads to unfavorable outcomes, including SC depletion, tissue degeneration, accumulation of oncogenic mutations and premature aging^5–7^. Recent insights about the SC quiescent state have allowed us to understand that rather it being a default phenomenon, SCs actively adopt this state to downregulate intrinsic fundamental processes such as cell cycle, cellular metabolism, and transcriptional output to withstand stress and genomic instability^8–11^. There are several regulatory cues that dictate SC quiescence or activation states; while extrinsic signals from the SC niche play an important role in dictating the transition between quiescent and active states, cell-intrinsic mechanisms need to respond to these environmental changes to facilitate this transition^12–14^. How extrinsic environmental changes influence intrinsic responses to regulate SC states remains a largely unanswered question.

The adult hair follicle (HF) serves as a powerful model to probe the regulatory mechanisms that modulate SC quiescence and activation. The HF undergoes cyclical bouts of regeneration, called the hair cycle, that can be divided into three successive phases of tissue remodeling – telogen (rest), anagen (growth) and catagen (regression)^15,16^. The long-lived hair follicle stem cells (HFSCs) that reside in a structurally defined “bulge” region of the HF are primarily quiescent but enter a brief period of activation during anagen onset to self-renew and generate progenitor cells that fuel hair regeneration^17–19^. However, the HFSC activation period is transient, and these cells quickly return to quiescence while the progenitor cells continue HF production. The quiescent and activation states of HFSCs are tightly governed by cell- intrinsic factors as well as extrinsic signals from the surrounding niche^20–23^. The temporal dynamics of HF regeneration offer a unique opportunity to dissect the mechanisms that establish and modulate the transcriptional programs governing HFSC quiescence and activation in response to niche signaling changes.

Histone modifications are plausible candidates for regulating HFSC states as they provide a higher- order control over gene expression^24,25^. Histone modifications, such as histone methylation, acetylation, or ubiquitination, modulate the overall structure of chromatin, determining the access of transcription factors (TFs) and other regulatory proteins to their target genes^26,27^. Thus, histone modifications can simultaneously regulate large networks of genes, giving them the potential to globally control SC states. Unlike TFs, which often act transiently^28,29^, histone modifications provide long-term stability but can rapidly remodel chromatin and regulate gene expression in response to changes in extrinsic cues^30,31^, therefore making them critical regulators of HFSC states, function, and identity over time.

### Histone mark H2AK119ub is dynamic during HFSC quiescence and activation

To identify if key histone modifications in control of gene expression are dynamic in HFSCs, we used immunofluorescence (IF) analysis coupled with the incorporation of 5-ethynyl-2’-deoxyuridine (EdU) to mark the quiescent and the earliest activated stages of the HF bulge region. We found no prominent changes in the levels of an active mark, tri-methylation of histone H3 lysine 4 (H3K4me3)^26,32^, between EdU- quiescent (P21 and P55) and EdU+ active HF bulges (P23 and P26) (**fig**. S1A). Compellingly, there was a distinct contrast in the distribution of the Polycomb-mediated histone modifications, which are known to co-repress target genes^33,34^. Notably, Polycomb repressive complex (PRC) 2-dependent tri-methylation of histone H3 lysine 27 (H3K27me3) remained steady between quiescent and active bulges (**fig**. S1B), whereas PRC1-dependent mono-ubiquitination of histone H2A lysine 119 (H2AK119ub) was diminished in active HF bulges in comparison to quiescent ones (**Fig.** 1A). We also conducted nuclear flow cytometry analysis of all three histone modifications in HFSCs identified as Sca1^-^, α6-integrin^high^ and CD34^+^ cells (**fig.** S1C). This assay verified that while there were no discernible changes in the levels of H3K4me3 and H3K27me3 in HFSCs of telogen and anagen skin (**fig.** S2A-B and **Fig.** 1C-D), the distribution of H2AK119ub+ HFSCs was significantly lower in activated bulges (∼63% in P23 and ∼53% in P26) when compared to quiescent ones (∼84% in P21 and ∼79% in P55) (**Fig.** 1B, E). IF analysis of H2AK119ub and HFSC stem cell marker, CD34^35^, confirmed that the level of H2AK119ub was downregulated in activated HFSCs of early-anagen skin (P23, P26) (**fig.** S2C). Collectively, these results identified that the repressive histone modification, H2AK119ub, oscillates between HFSC quiescent and active states.

**Figure 1.**
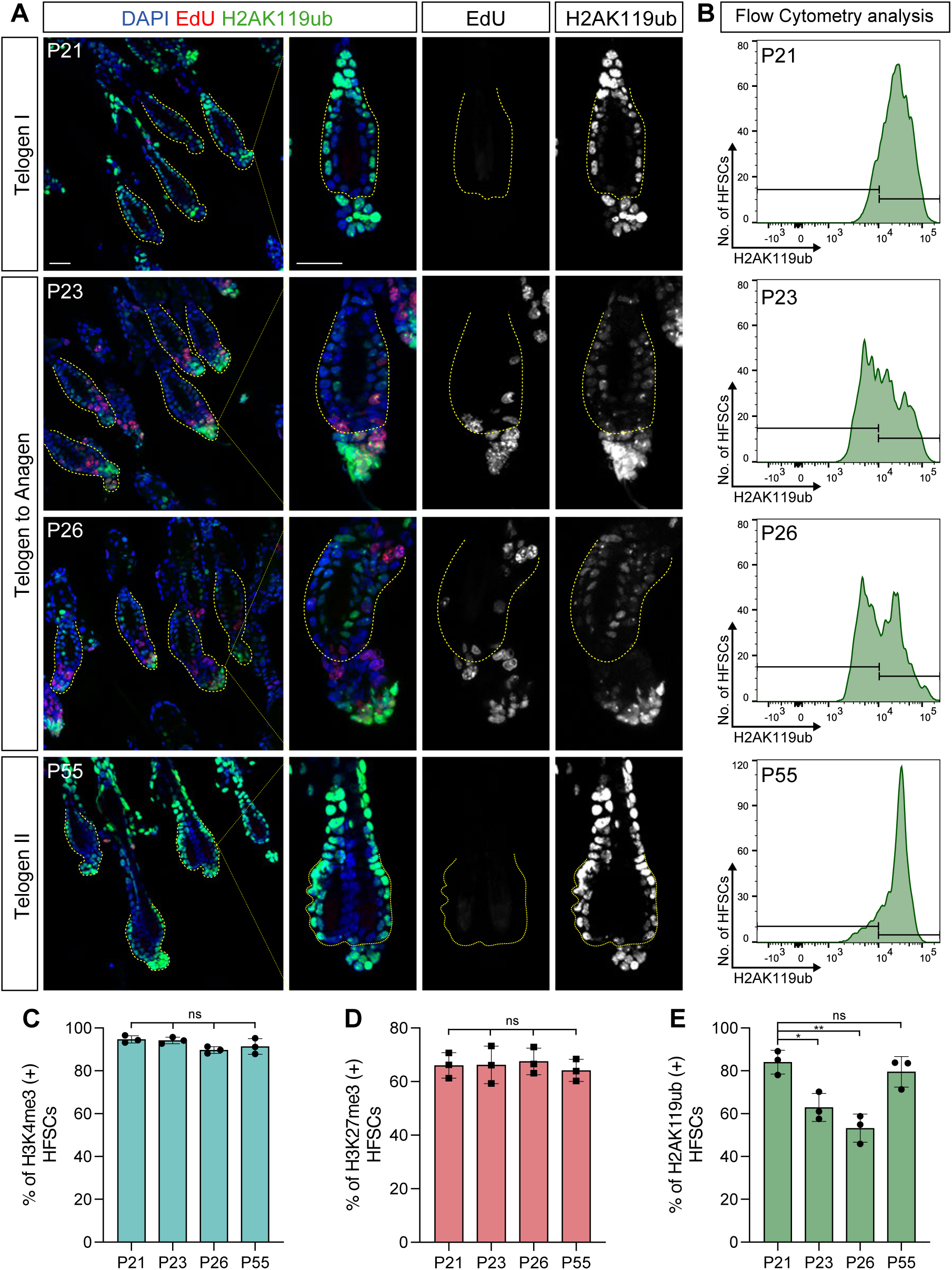
H2AK119ub levels are dynamic between quiescent and activated HFSCs. (**A**) Epidermal whole-mount IF analysis of H2AK119ub (green), EdU (red), and DAPI (blue) in skin samples from telogen I (P21), early anagen (P23, P26), and telogen II (P55) showing that EdU+ proliferating bulges of P23 and P26 skin have markedly lower levels of H2AK119ub when compared to EdU- quiescent bulges of P21 and P55 skin. EdU and H2AK119ub channels shown in grayscale. The bulge region is indicated with a yellow dashed line. (**B**) Nuclear flow cytometry histograms showing the intensity of H2AK119ub in HFSCs isolated from dorsal skin of P21, P23, P26, and P55 animals. (**C-E**) Bar graphs representing the percentage of H3K4me3(+), H3K27me3(+), and H2AK119ub(+) HFSCs detected using flow cytometry, respectively (n = 3 animals from independent litters per time point). One-way ANOVA (Dunnett’s multiple comparisons test) was conducted. P-values for (**C**) = 0.9861 (P21 vs. P23), 0.700 (P21 vs. P26), and 0.2618 (P21 vs. P55); P-values for (**D**) = 0.9999 (P21 vs. P23), 0.9714 (P21 vs. P26), and 0.9480 (P21 vs. P55); P-values for (**E**) = 0.0103 (P21 vs. P23), 0.0010 (P21 vs. P26), and 0.7395 (P21 vs. P55). Scale bar: 25μm.

### H2AK119ub preserves HFSC quiescence in the adult skin

To determine whether the dynamic changes in H2AK119ub levels observed in quiescent and activated HFSCs have a functional role in regulating HFSC states, we generated conditional-inducible PRC1-null mice by crossing *K15-CrePR*^36^ mice with *Ring1a*-null *Ring1b^flox/flox^* mice (*K15-CrePR: Ring1a^-/-^, Ring1b^flox/flox^* = PRC1 i2KO) (**fig.** S3A). We topically applied RU-486 to the shaved back skin of PRC1 i2KO mice during telogen II to delete the essential E3 catalytic core of PRC1 complex, RING1B, and confirmed the efficient ablation of its corresponding histone mark H2AK119ub, specifically in the HF bulge, including the HFSCs (**fig.** S3B-D). IF analysis of H3K27me3 confirmed that its levels remain unchanged upon the loss of PRC1 function (**fig.** S3E), thereby establishing that H2AK119ub distribution in the bulge is not necessary to facilitate the deposition of the PRC2-dependent H3K27me3 histone mark. PRC1 is a potent regulator of SC fate and represses pro-apoptotic genes to conserve SC function in developing and homeostatic tissues^37–39^. However, IF analysis of control and PRC1 i2KO skin post-induction revealed that loss of PRC1 neither alters HFSC fate as expression of HFSC fate marker, SOX9^40^, is comparable between control and PRC1-null HFSCs (**fig.** S3F) nor does it lead to the expression of pro-apoptotic marker P19 (ARF) (**fig.** S3G) or programmed cell death marker activated-Caspase 3 (Ac-CASP3) (**fig.** S3H).

Hematoxylin and Eosin (H&E) analysis of PRC1 i2KO and control littermate skins revealed that while control HFs remained in telogen, the PRC1 i2KO HFs underwent premature anagen and recovered their hair coat significantly faster than control littermates (**fig.** S4A and **Fig.** 2A-C). To monitor the progression through subsequent anagen and catagen phases of the hair cycle, we shaved the recovered hair coat in PRC1 i2KO mice and observed telogen skin, characterized by its pink hue, confirming the unperturbed completion of the hair cycle (**fig.** S4B). Interestingly, PRC1 i2KO mice initiated the next anagen phase and recovered their second hair coat post-induction, while the control littermates were yet to recover their first one (**fig.** S4B). As PRC1 i2KO mice undergo precocious hair cycling, we tested if loss of H2AK119ub leads to premature activation of the HF bulge. Analysis of proliferation by EdU incorporation showed no significant proliferative events in either control or PRC1 i2KO HFs at P70 (**fig.** S4C, **Fig.** 2E); however, by P73 ∼53% of the HFs in PRC1 i2KO were EdU+, a percentage that increased to around 72% in P75 samples (**Fig.** 2D-E). To complement our *in vivo* observation of precocious HFSC activation in PRC1 i2KO mice, we implemented a previously described fluorescent activated cell sorting (FACS) strategy^41,42^ for isolating quiescent HFSCs as EpCAM^+^, Sca1^-^, α6-integrin^high^, CD34^+^ cells from P70 control and PRC1 i2KO mice (**fig.** S4D) and cultured them on fibroblast feeders. Indeed, PRC1 i2KO HFSCs formed significantly bigger colonies, indicating that loss of H2AK119ub in quiescent HFSCs leads to an increased proliferative potential (**Fig.** 2F).

**Figure 2.**
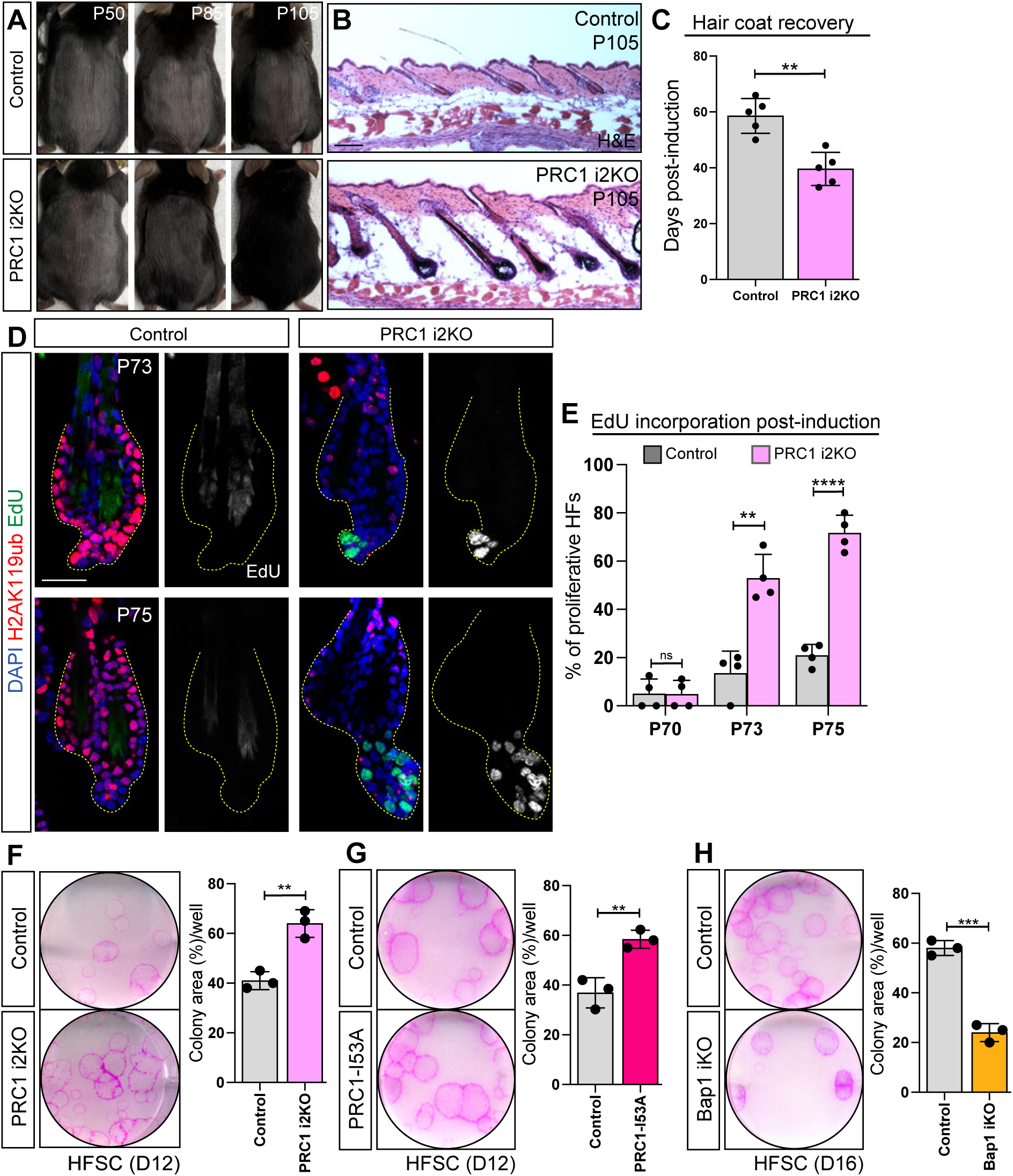
H2AK119ub preserves HFSC quiescence. (**A**) Images of back skin of control and PRC1 i2KO mice at start of induction (P50), 25 days (P85), and 45 days (P105) after induction treatment, respectively. (B) H&E analysis of skin sections from P105 control and PRC1 i2KO back skin. (**C**) Bar graph showing that PRC1 i2KO mice recover hair coat faster than control littermates (n = 5 animals from at least three different litters). A non-parametric Mann-Whitney t-test was conducted. P-value = 0.0079. (**D**) Epidermal whole-mount IF analysis of H2AK119ub (red), EdU (green) and DAPI (blue) in the HF bulge region of of P73 and P75 control and PRC1 i2KO mice. EdU channel shown in grayscale. (**E**) A bar graph showing the percentage of EdU+ HFs in P70, P73 and P75 control and PRC1 i2KO mice (n= 50-60 HFs/animal from 4 independent biological replicates for each group). An unpaired t-test was conducted. P-value = 0.9591 (P70); 0.0010 (P73); 0.00002 (P75). (**F**) HFSC colonies from P70 control and PRC1 i2KO mice. Bar graph depicting PRC1-null HFSCs made bigger colonies compared to control (n = 3 animals from at least two independent litters). An unpaired t-test was conducted. P-value = 0.0063. (**G**) HFSC colonies from P70 control and PRC1-I53A mice. Bar graph depicting I53A mutant HFSCs made bigger colonies compared to control (n = 3 animals from at least two independent litters). An unpaired t-test was conducted. P-value = 0.0037. (**H**) HFSC colonies from P70 control and Bap1 iKO mice. Bar graph showing Bap1-null HFSCs made smaller colonies compared to control (n = 3 animals from at least two independent litters). An unpaired t-test was conducted. P-value = 0.0037. Scale bar for IF images: 25μm; for H&E images: 50μm.

To discriminate if the phenotype observed in PRC1 i2KO mice was predominantly dependent on the loss of PRC1’s catalytic activity of depositing mono-ubiquitination^43^, we generated conditional-inducible PRC1 catalytic-inactive mice (*K15-CrePR: Ring1a^-/-^,Ring1b^I53A/flox^* = PRC1-I53A), in which the *Ring1b* gene carries an I53A point mutation that abolishes its E3-ligase activity and the establishment of H2AK119ub in HFSCs while maintaining all other PRC1 functions^44^ (**fig.** S5A-B). Matching our observations with PRC1 i2KO mice, PRC1-I53A mice also entered anagen prematurely and recovered their hair coat sooner than control littermates (**fig.** S5C). *In vitro*, quiescent PRC1-I53A HFSCs also made bigger colonies compared to control, phenocopying PRC1 i2KO HFSCs (**Fig.** 2G).

Finally, we asked if inhibiting the dynamic regulation of H2AK119ub would impact HF regeneration and HFSC proliferative status. BAP1 is a ubiquitin hydrolase that works antagonistically to PRC1 to remove the mono-ubiquitination of histone H2A lysine 119^45^. We generated conditional-inducible *Bap1*-null mice (*K15-CrePR: Bap1^flox/flox^*= Bap1 iKO) and confirmed the downregulation of *Bap1* RNA in HFSCs by conducting RT-qPCR (**fig.** S5D-E). Remarkably, loss of Bap1 in the HFSCs not only resulted in a delayed anagen entry compared to control littermates (**fig.** S5F), but the proliferative potential of Bap1-null HFSCs was also significantly compromised (**Fig.** 2H), which completely opposed our observations with PRC1 i2KO and PRC1-I53A HFSCs. Overall, these genetic studies highlight that H2AK119ub is not only the primary regulator of HFSC quiescence, but the temporal modulation of H2AK119ub in HFSCs is essential for the timely transition of HFSC quiescence to activation to fuel HF regeneration.

### Dysregulation of quiescence upon loss of H2AK119ub results in HFSC exhaustion

Unchecked proliferation and loss of quiescence has been implicated in SC aging of various adult tissues^6,46^; therefore, we asked if prolonged loss of H2AK119ub and enhanced proliferative potential of PRC1-null HFSCs is detrimental to HFSC longevity. Intriguingly, 10-month old (10M) PRC1 i2KO mice presented a thinning hair coat when compared to 10M control littermates – a landmark of HF aging (**Fig.** 3A). IF analysis showed that control HFs had uniform CD34 expression and had two or more bulges, indicative of animals undergoing at least two hair cycles. Comparatively, CD34 expression in PRC1-null HFSCs was grossly downregulated and ∼51% of the bulges exhibited defective morphology (**Fig.** 3B-C). Flow cytometry analysis showed that 10M PRC1 i2KO mice had a significant reduction in their HFSC pool compared to corresponding controls, a phenotype not observed in 2M animals (**Fig.** 3D, **fig.** S5G). Next, we tested if the remaining HFSCs in 10M PRC1 i2KO mice sustained their enhanced proliferative potential as observed in animals 10 days post-induction (**Fig.** 2F). When cultured on fibroblast feeder cells,10M PRC1 i2KO HFSCs made significantly smaller colonies than 10M control HFSCs (**Fig.** 3E). Together, these results indicate that H2AK119ub mediated HFSC quiescence is indispensable for conserving HFSC function and longevity.

**Figure 3.**
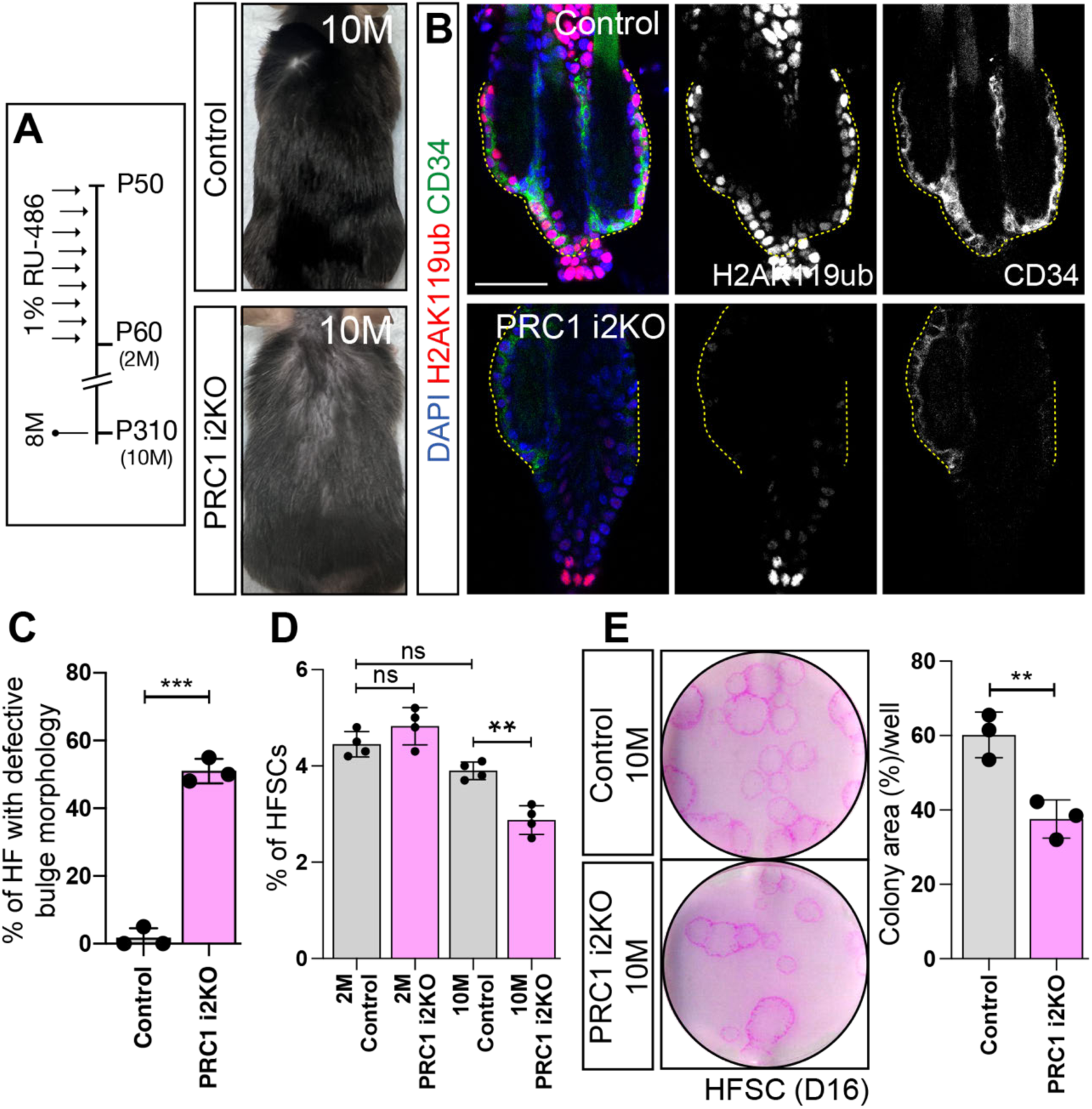
Loss of H2AK119ub in the bulge leads to HFSC exhaustion over time. (**A**) Schematic showing the experimental strategy for long-term observation of control and PRC1 i2KO mice, along with representative images of back skin from control and PRC1 i2KO mice 8 months post-induction (**B**) Epidermal whole-mount IF analysis of H2AK119ub (red), CD34 (green), and DAPI (blue) in HF bulges of 10M control and PRC1 i2KO mice. H2AK119ub and CD34 channels shown in grayscale. (**C**) A bar graph showing that 51% of PRC1 i2KO bulges were defective in morphology and CD34 expression (n = 3 animals from at least two independent litters). An unpaired t-test was conducted. P-value = 0.0001. (**D**) A bar graph showing flow cytometry analysis of the percentage of HFSCs in 2M control vs. PRC1 i2KO and 10M control vs. PRC1 i2KO animals (n = 4 animals from at least two independent litters). A one-way ANOVA (Tukey’s multiple comparison’s test) was conducted. P-value = 0.3136 (2M control vs. 2M PRC1 i2KO), 0.0845 (2M control vs. 10M control), and 0.0016 (10M control vs. 10M PRC1 i2KO). (**E**) HFSC colonies from 10M control and PRC1 i2KO mice. Bar graph showing 10M PRC1-null HFSCs made smaller colonies compared to control (n = 3 animals from at least two independent litters). An unpaired t-test was conducted. P-value = 0.0080. Scale bar: 25μm.

### H2AK119ub directly represses HFSC proliferation program in quiescent HFSCs

To identify the processes de-regulated in quiescent HFSCs upon the loss of H2AK119ub, we purified HFSCs from the back skin of P70 control and PRC1 i2KO mice (**fig.** S4D), prior to the onset of proliferation (**fig.** S4C), and conducted RNA sequencing (RNA-seq). Differential gene expression analysis revealed significant downregulation of 176 genes and upregulation of 774 genes in PRC1-null HFSCs (**Fig.** 4A). As H2AK119ub is primarily involved in repressing gene expression, we focused on identifying the processes upregulated upon its ablation. Gene ontology (GO) analysis revealed that genes upregulated in P70 PRC1-null HFSCs were associated with proliferation promoting processes such as cell cycle and cell division (**fig.** S6A). Inversely, expression of known HFSC quiescence maintaining factors including *Nfatc1*, *Foxc1* and *Bmp6*^47–50^ remained unchanged (**fig.** S6B), indicating that H2AK119ub regulates quiescence independent of these factors. We also aimed to identify if loss of PRC1 leads to the induction of an activated HFSC state. Indeed, gene-set enrichment analysis (GSEA) with a previously described anagen-enriched gene data set^42^ revealed that PRC1-null HFSCs are similar to activated HFSCs, with upregulation of 166- anagen enriched genes (**Fig.** 4B). GO analysis of these genes revealed an overwhelming association with cell cycle, cell division, mitosis, and proliferation processes (**Fig.** 4C). Among these genes were, epidermal SC activation regulators such as *Ptch2*^51^, and *MycN*^52^ and cell cycle and cell division genes such as *Ccna2*^53^, *Ccnb2*^54^, *Cdc7*^55^, *Cdk1*^56^, *Mki67*^57^, *Kif23*^58^ and *Top2a*^59^ (**Fig.** 4C). Because writing and erasure of histone modification is a labile process^60^ and robust changes in gene expression dependent on the removal of histone modifications may take a longer period, we also conducted RNA-seq with HFSCs isolated from the back skin of P75 control and PRC1 i2KO mice. Differential gene expression analysis revealed even larger changes in PRC1-null HFSCs with 1546 upregulated genes (**fig.** S6C). Unsurprisingly, GO analysis revealed that top biological processes associated with P75 upregulated genes were associated with cell cycle and cell division (**fig.** S6D). Merging with anagen-enriched data set revealed 310 genes upregulated in P75 PRC1 i2KO HFSCs were anagen-enriched (**fig.** S6E). Together, these transcriptional studies revealed that loss of H2AK119ub in HFSCs leads to a progressive establishment of a transcriptional landscape that is akin to activated HFSCs.

**Figure 4.**
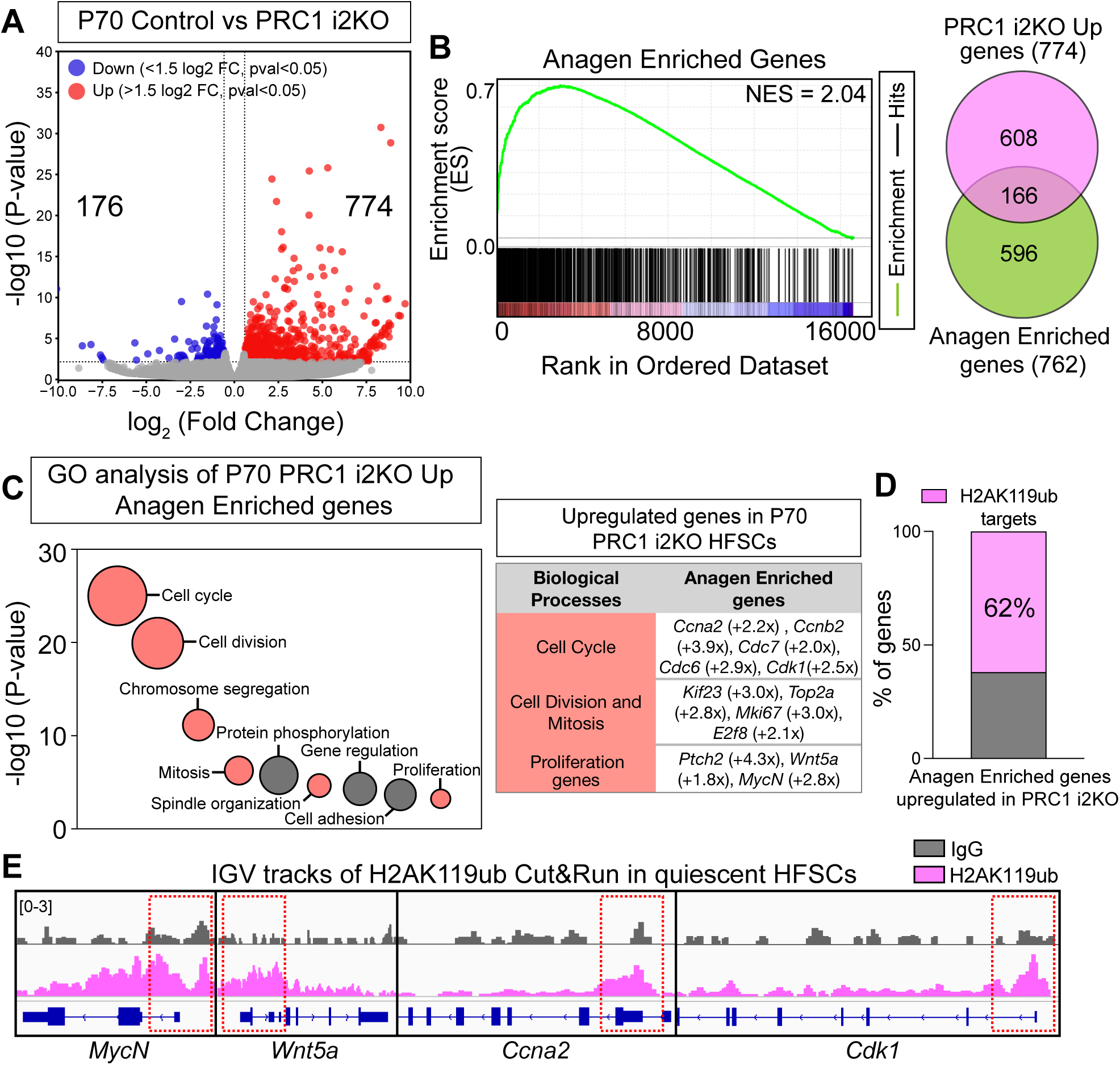
Loss of H2AK119ub initiates a proliferation promoting transcriptional landscape in HFSCs. (**A**) Volcano plot showing the differentially expressed (DE) genes in FACS-purified HFSCs from P70 control and PRC1 i2KO mice. Genes with absolute fold change ≥ 1.5 and adjusted p-value < 0.05 were considered significantly upregulated or downregulated for further analysis. RNA-seq analysis was done on three biological replicates for each group from at least two independent litters. (**B**) Gene set enrichment analysis of genes differentially expressed in PRC1 i2KO HFSCs with anagen enriched genes (NES = 2.04, P-value <1E^-10^) and Venn diagram showing 166 anagen enriched genes were upregulated in PRC1 i2KO HFSCs. (**C**) Gene Ontology (GO) analysis of 166 anagen enriched genes upregulated in PRC1 i2KO HFSCs and table listing the fold change expression of selected DE genes. (**D**) Stacked graph showing 62% of anagen enriched genes upregulated in PRC1 i2KO HFSCs are H2AK119ub targets. (**E**) IGV tracks showing H2AK119ub occupancy on proliferation promoting and cell cycle genes in quiescent HFSCs.

Next, we asked if H2AK119ub directly represses these anagen-enriched genes in quiescent HFSCs. Hence, we conducted Cleavage Under Targets & Release Using Nuclease (Cut&Run) with antibodies against H2AK119ub in quiescent HFSCs isolated from the back skin of telogen II (P55) C57BL/6 wildtype mice. Analysis of H2AK119ub occupancy revealed that 68.7% of genes upregulated in P70 and 52.5% upregulated in P75 PRC1 i2KO HFSCs were H2AK119ub targets (p < 0.05, Fisher’s test) (**fig.** S6F- G). Specifically, 62% of anagen-enriched genes upregulated in P70 PRC1-null HFSCs were demarcated by H2AK119ub in quiescent HFSCs (**Fig.** 4D). Notably, proliferation-promoting genes, *Wnt5a*^61^*, Ptch2*, *MycN*, and key cell cycle genes, *Ccna2, Cdk1, Cdc7* and *Kif23*, were direct targets of H2AK119ub- mediated repression (**Fig**. 4E, **fig.** S6H) indicating that H2AK119ub directly represses the HFSC activation promoting transcriptional landscape to preserve HFSC quiescence.

### Quiescent promoting FGF niche signaling maintains H2AK119ub levels in quiescent HFSCs

Our data show that H2AK119ub directly represses key cell cycle and proliferation promoting genes to safeguard quiescence in HFSCs. This repression, however, needs to be alleviated during HFSC activation. Since the level of H2AK119ub decreases in anagen HFSC (**Fig.** 1A-B, E), we aimed to identify if extrinsic niche signaling in the HF bulge regulates H2AK119ub levels during telogen to anagen transition. Several signaling pathways in the HF bulge niche are dynamic during the transition of HFSC quiescence to activation, including the fibroblast growth factor (FGF) signaling^62,48,22^. During telogen, the Keratin (Krt) 6+ non-proliferative keratinocytes located adjacent to the HFSCs serves as a niche and produces high levels of inhibitory FGF18 ligand which is downregulated prior to HFSC activation^48,62^ (**Fig.** 5A, top); thereby highlighting the dynamic nature of this signaling pathway in maintaining HFSC quiescence. Notably, removal of the Krt6+ inner bulge cells, via depilation, during telogen results in precocious activation of HFSCs^48^ and conditional knock-out of FGF18 in the epidermis, including the Krt6+ cells, leads to precocious anagen entry and rapid successive hair cycling^63^. Based on these previous findings, we first tested if removal of FGF18 producing Krt6+ bulge cells via depilation impacts H2AK119ub levels in HFSCs. Therefore, we depilated the telogen skin of P60 C57BL/6 wildtype mice and assayed for EdU incorporation and H2AK119ub levels. We found that compared to the quiescent bulges of control P62 skin which maintained H2AK119ub levels with no EdU incorporation, depilation led to drastic downregulation of H2AK119ub levels in addition to bulge proliferation, signified by prominent EdU+ cells (**fig.** S7A).

**Figure 5.**
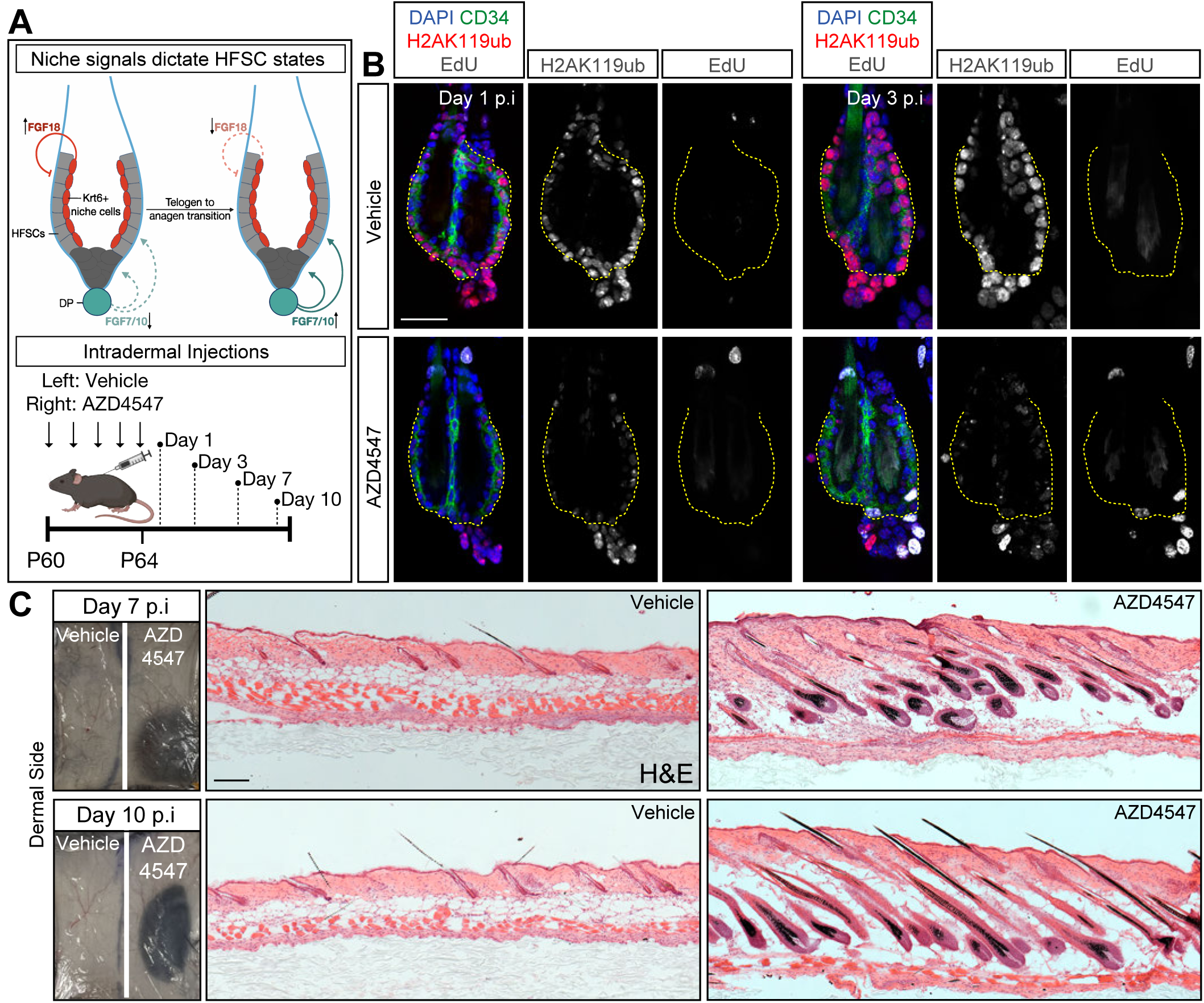
H2AK119ub levels diminish upon inhibition of niche FGF signal. (**A**, top) Schematic of FGF signaling pathways that mediate HFSC quiescence and activation. (**A**, bottom) Schematic of FGF inhibition experimental strategy by intradermal injections. (**B**) Epidermal whole-mount IF analysis of H2AK119ub (red), CD34 (green), EdU (gray), and DAPI (blue) in skin samples from vehicle and AZD4547 injected site 1 day and 3 days post-injection (p.i) regimen show that H2AK119ub levels reduce upon AZD4547 injections 1 day post-injection and HF bulges start proliferating within 3 days post-injection. H2AK119ub and EdU channels shown in grayscale (**C**) H&E analysis of skin sections from vehicle and AZD4547 injected site 7 days and 10 days post-injection regimen. Scale bar for IF images: 25μm; for H&E images: 100μm.

Next, we asked if pharmacological inhibition of the endogenous FGF signaling, by targeting fibroblast growth factor receptors (FGFR), in the skin would lead to downregulation of H2AK119ub, induce HFSC activation and precocious anagen entry. Therefore, we conducted intradermal injections in C57BL/6 wildtype mice with vehicle (PBS+DMSO) and a selective FGFR 1-3 inhibitor, AZD4547, over the course of 5 days during mid-telogen (P60) when the FGF18 ligands are high and FGF7/10 ligand are very low^62^ (**Fig.** 5A), thereby targeting the inhibitory FGF18 pathway. We then assayed for H2AK119ub levels and proliferation in the bulges of the vehicle and AZD4547 injected site 1 day after completion of the injection regimen. Remarkably, compared to HFSCs in the vehicle samples, H2AK119ub levels were drastically downregulated in the HFSCs of the AZD4547 injected site (**Fig.** 5B, left), albeit no proliferation was detected. However, we observed that the bulges in AZD4547 injected site started proliferating 3 days post- injection regimen, while the vehicle site remained quiescent (**Fig.** 5B, right). By day 7 post-injection, the AZD 4547 site had visible darkening of the dermis, indicating active anagen phase, which was even more prominent by day 10 post-injection regimen (**Fig.** 5C, left). H&E staining of skin sections from these time- points verified that while the HFs in the vehicle injected site remained in telogen, the HFs in the AZD 4547 injected site were in the anagen phase of the hair cycle (**Fig.** 5C, right). Subsequently, by day 14, we observed hair coat recovery specifically at inhibitor injected site (**fig.** S7B). To eliminate that inhibition of FGF18 signaling affected any other histone modifications and this phenomenon was unique to H2AK119ub, we assayed for H3K27me3 and H3K4me3 levels in bulges of vehicle and AZD4547 injected site and confirmed that levels of these histone modifications remained unaffected by AZD4547 (**fig.** S7C- D). Together, these studies unveiled that the quiescent maintaining FGF signaling from the bulge niche was not only regulating H2AK119ub levels in HFSCs to sustain quiescence but downregulation of H2AK119ub via FGF signaling inhibition was sufficient to induce precocious HFSC activation and anagen entry.

## Discussion

Dynamic changes in histone modifications have been linked to genomic instability, cellular stress, senescence, aging, and progression of cancer^64,65^. However, our knowledge regarding the dynamic regulation of histone modifications in dictating SC states has been limited. The regulation of HFSC activation is remarkably transient and both temporal and spatial regulation of cell-intrinsic and extrinsic factors tightly orchestrate the transition of HFSC quiescence to activation^21–23^. Whether dynamic gene regulatory mechanisms mediated by histone modifications functionally contribute to this transient process was unclear. In this study, we have successfully identified that a repressive histone modification, H2AK119ub, is uniquely dynamic between quiescent and activated HFSCs. Specifically, H2AK119ub levels are maintained during quiescence, but there is a global decrease of its levels in the earliest stages of HFSC activation, contrary to our observations with H3K4me3 and H3K27me3 histone marks that have been described as guardians of cellular identity and plasticity^41,66,67^. Our functional studies probing the importance of this phenomenon during physiological hair cycling revealed that H2AK119ub is essential for maintaining HFSC quiescence. Specifically, ablation of this repressive mark leads to increased HFSC proliferative potential, premature HFSC activation, and untimely onset of HF regeneration. Conversely, inhibiting the decrease of H2AK119ub delays hair cycling and compromises its proliferative potential *in vitro*. Therefore, H2AK119ub has emerged not only as an indispensable cell-intrinsic regulator of a SC population, but its temporal regulation is also essential for timely regeneration.

Maintaining quiescence has been postulated to be critical for preserving SC numbers and regenerative capacity long-term^6,8,11^. Studies focused on dissecting the role of TFs, such as NFATc1 and FOXC1, in mediating HFSC quiescence had revealed that loss of these factors, leads to premature SC exhaustion, HF miniaturization and compromised regenerative potential^47,49,68,69^ – all hallmarks of aging. Our investigations speculating the long-term requirement of H2AK119ub-mediated quiescence mirror these previous observations. Sustained loss of H2AK119ub in HFSCs results in premature thinning of hair coat, defective HF bulge morphology, dramatic reduction in HFSC pool, and reduced proliferative capacity *in vitro*. Hence, highlighting that dysregulation of quiescence maintaining processes under H2AK119ub regulation leads to the premature onset of several aging-related phenotypes. Importantly, as NFATc1, TCF3/4 and BMP6, are not dysregulated in H2AK119ub-null quiescent HFSCs, our studies identify an alternative quiescence promoting instructive mechanism mediated by a histone modification. Although initial loss of H2AK119ub did not affect HFSC function and differentiation indicated by proper progression through hair cycle, the premature onset of several aging-associated phenotypes highlights H2AK119ub’s role as a guardian of HFSC longevity. If H2AK119ub plays a role in other cell types of the HF to facilitate multiple rounds of regeneration, as implied by studies conducted in a depilation model^70^, remains to be investigated.

Self-renewal and differentiation of stem cells are fundamentally associated with progression through cell cycle to ensure proper tissue regeneration^71^. While timely initiation of a proliferative program in stem cells is vital for regeneration, these processes must be strictly regulated to conserve SC quiescence and longevity. Not surprisingly, during HFSC activation, several cell cycle and cell division related genes, along with proliferation promoting signaling pathways, are upregulated in comparison to quiescent HFSCs. Additionally, loss-of-function of quiescence maintaining TFs factors in HFSCs also leads to an upregulation of cell cycle genes^47,50^. Although the establishment of a proliferation promoting transcriptional landscape is one of the earliest intrinsic responses of HFSC activation, our knowledge of their direct regulation during the transition from quiescence to activation has been limited. Our transcriptional and chromatin-mapping studies revealed that key HFSC proliferation promoting, cell cycle, and cell-division genes were direct targets of H2AK119ub-mediated repression in quiescent HFSCs, thereby providing a global view of how a chromatin regulatory process directly regulates proliferation to preserve HFSC quiescence. It is worth noting that H2AK119ub directly represses only a subset of HFSC proliferation promoting genes, but we infer that alleviation of H2AK119ub-mediated repression on these genes results in a domino effect, thereby inducing the establishment of transcriptional landscape that promotes HFSC activation and subsequent HF regeneration.

Concomitant with cell-intrinsic factors, HFSC quiescent and activated states are also regulated by the temporal and spatial dynamics of extrinsic signaling pathways. Especially, BMP and FGF signaling pathways in the HF microenvironment have profound roles in governing HFSC states. While inhibitory BMP6 and FGF18 signals from the Krt6+ inner bulge cells promote HFSC quiescence, their levels diminish prior to HFSC activation and increasing levels of FGF7/10 signaling from the dermal papilla (DP) promotes HF regeneration^48,62^. While the temporal expression of these signaling pathways has been implicated in modulating the expression of cell-intrinsic quiescence factors^47^, their role in remodeling the epigenetic landscape to control HFSC quiescence was previously unknown. Our pharmacological studies targeting the inhibitory FGF signaling pathway not only led to the selective reduction in H2AK119ub levels in HFSCs, but it was sufficient to induce precocious proliferation and premature anagen induction, similar to the PRC1 i2KO animals. Together, these studies provide us an insight into the signaling-epigenetic axis in control of HFSC states.

In conclusion, our study showcases that the global levels of a repressive histone modification, H2AK119ub, oscillates in HFSCs in response to extrinsic niche signaling to control the induction of a proliferative transcriptional landscape to mediate the transition from HFSC quiescence to activation and preserve HFSC longevity. Our findings open the door to future investigations into identifying if this signaling-epigenetic axis regulates stem cell quiescence in other organs, as well as allow us to explore if dysregulation of this regulatory process leads to unchecked cellular proliferation and loss of quiescence that is often associated with malignant transformation of cells.

---

**Supplemental Figure 1.**
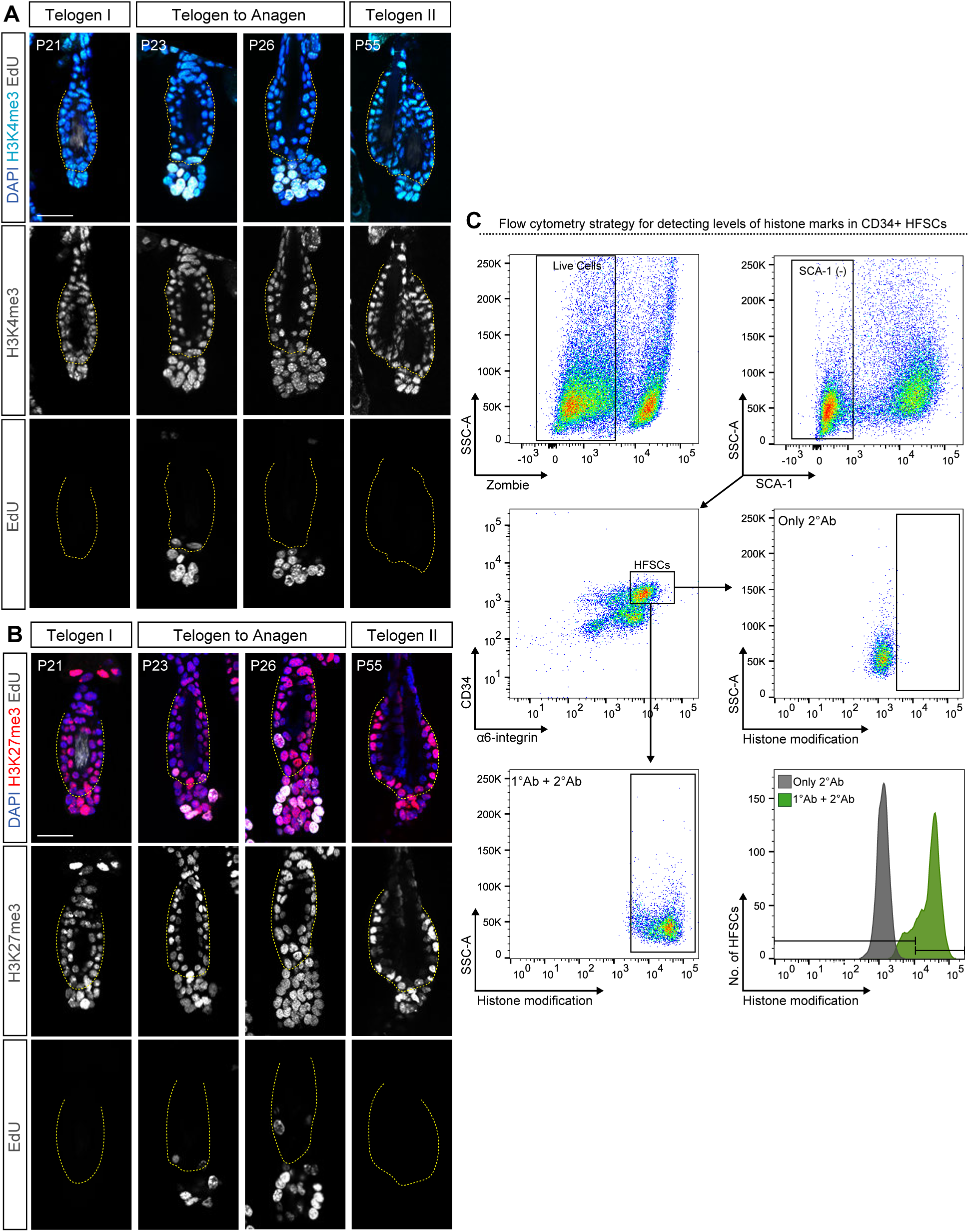
(Related to Fig. 1) Levels of H3K4me3 and H3K27me3 are maintained in quiescent and activated HF bulges. (**A**) Epidermal whole-mount IF analysis of H3K4me3 (cyan), EdU (gray), and DAPI (blue) in skin samples from telogen I (P21), early anagen (P23, P26), and telogen II (P55) showing no discernible differences in the levels of H3K4me3 between EdU^+^ proliferating bulges of P23 and P26 and EdU^-^ quiescent bulges of P21 and P23 skin. EdU and H3K4me3 channels shown in grayscale. The bulge region is indicated with a yellow dashed line. (**B**) Epidermal whole-mount IF analysis of H3K27me3 (red), EdU (gray), and DAPI (blue) in skin samples from telogen I (P21), early anagen (P23, P26), and telogen II (P55) showing no discernible differences in the levels of H3K27me3 between EdU^+^ proliferating bulges of P23 and P26 and EdU^-^ quiescent bulges of P21 and P23 skin. EdU and H3K27me3 channels shown in grayscale. The bulge region is indicated with a yellow dashed line. (**C**) Flow cytometry strategy of detecting histone modification levels in HFSCs identified as Sca1^-^, α6-integrin^high^, and CD34^+^. Cells stained with concentration matched secondary (2°) antibody served as a negative control. Histograms were generated to identify the number of HFSCs that were positive for nuclear staining of histone modifications. Scale bar: 25μm.

**Supplemental Figure 2.**
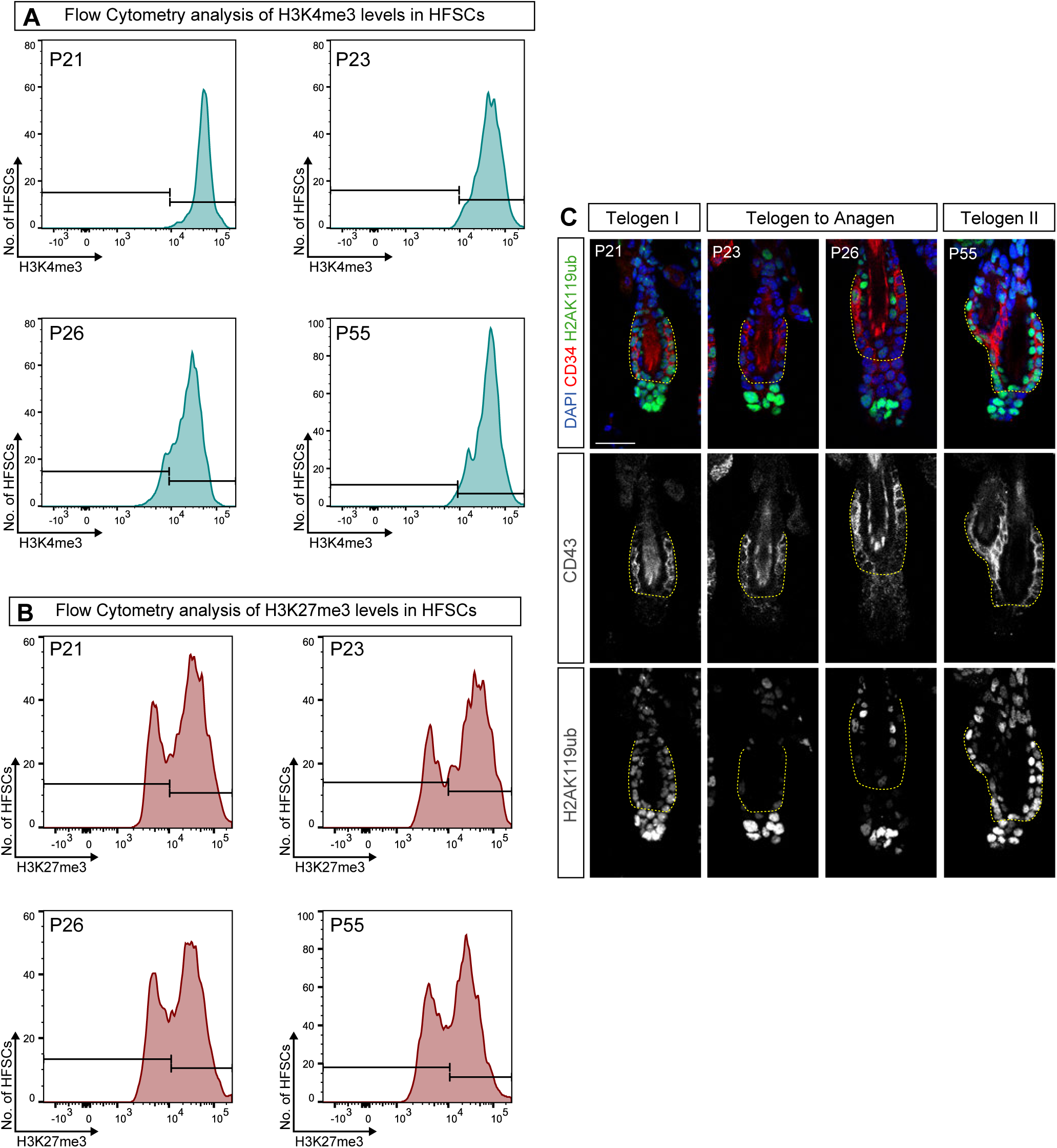
(Related to Fig. 1) Levels of H3K4me3 and H3K27me3 are maintained in quiescent and activated HF bulges. (**A** - **B**) Nuclear flow cytometry histograms depicting the intensity of H3K4me3 and H3K27me3 in HFSCs isolated from dorsal skin of P21, P23, P26, and P55 animals, respectively. (**C**) Epidermal whole-mount IF analysis of skin samples from mice in telogen I (P21), early anagen (P23, P26), and telogen II (P55) stained with CD34 (red), H2AK119ub (green), and DAPI (blue) showing that HFSCs of activated P23 and P26 bulges downregulate H2AK119ub compared to the HFSCs of quiescent P21 and P55 bulges. Scale bar: 25μm.

**Supplemental Figure 3.**
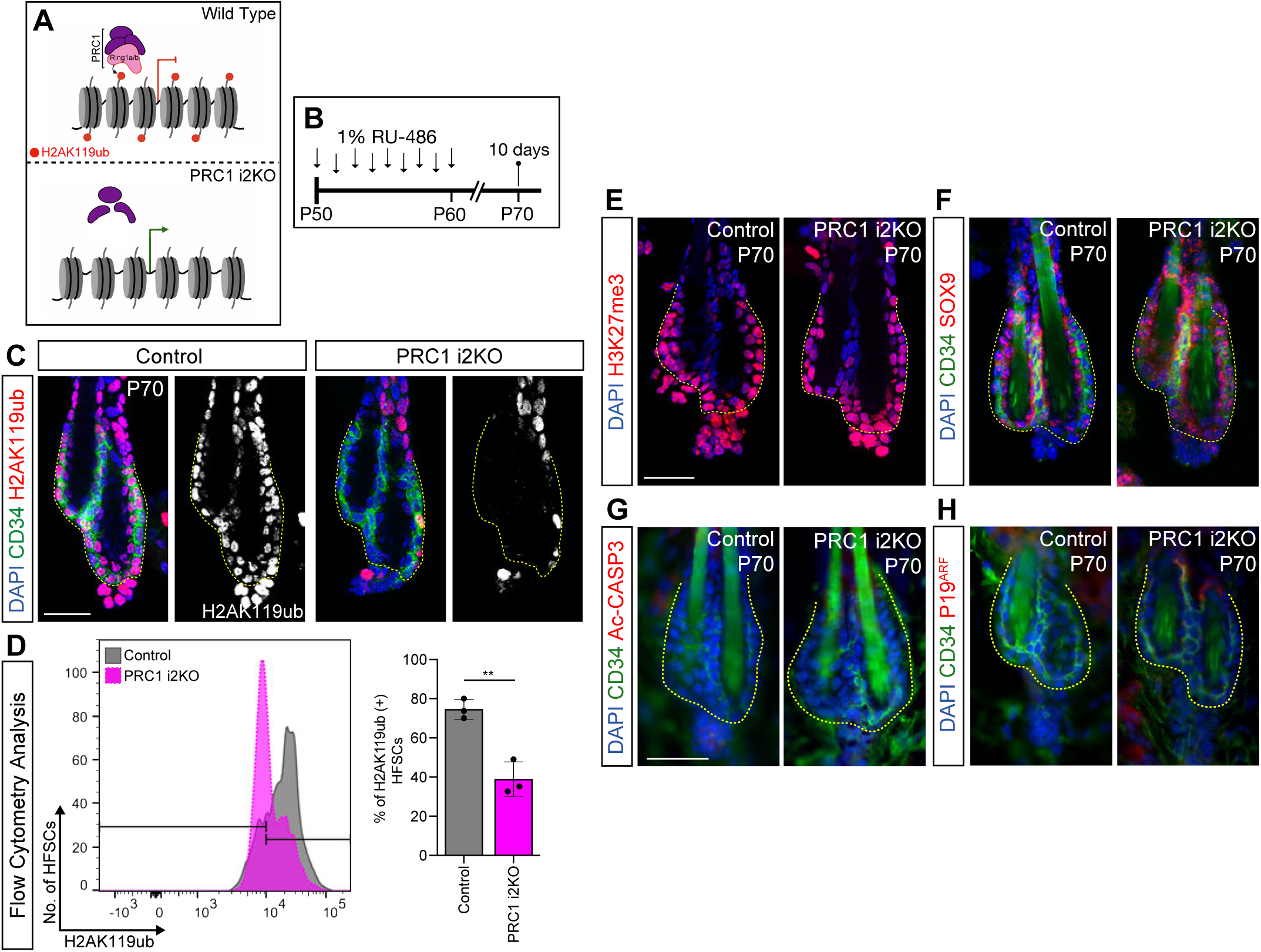
(Related to Fig. 2) Efficiency of H2AK119ub ablation in the bulge using *K15- CrePR*. (**A**) Schematic illustrating that both RING1A and RING1B function are ablated in PRC1 i2KO mice resulting in ablation of H2AK119ub. (**B**) Schematic showing the experimental strategy to induce *K15- CrePR* activity in telogen II (P50) HFs and skin samples were collected at P70 for analyzing ablation efficiency. (**C**) Epidermal whole-mount IF analysis of H2AK119ub (red), CD34 (green), and DAPI (blue) in HFSCs of P70 control and PRC1 i2KO HF bulges. H2AK119ub channel shown in grayscale. (**D**) Nuclear flow cytometry histograms and corresponding bar graph showing that compared to control, number of H2AK119ub(+) HFSCs are significantly reduced in PRC1 i2KO mice post-induction (n = 3 animals from at least two independent litters). An unpaired t-test was conducted. P-value = 0.0037. (**E**) Epidermal whole-mount IF analysis of H3K27me3 (red) and DAPI (blue) show no changes in the levels of these histone modifications between control and PRC1 i2KO HF bulges. (**F**) Epidermal whole-mount IF analysis SOX9 (red), CD34 (green), and DAPI (blue) show no changes in SOX9 expression between control and PRC1 i2KO HF bulges. (**G**) IF analysis of activated-CASPASE3 (red), CD34 (green), and DAPI (blue) in P70 control and PRC1 i2KO skin sections showing alation of H2AK119ub does not lead to cell death. (**H**) IF analysis of P19^ARF^ (red), CD34 (green), and DAPI (blue) in P70 control and PRC1 i2KO skin sections showing no differences in the levels of P19^ARF^ expression. Scale bar: 25μm.

**Supplemental Figure 4.**
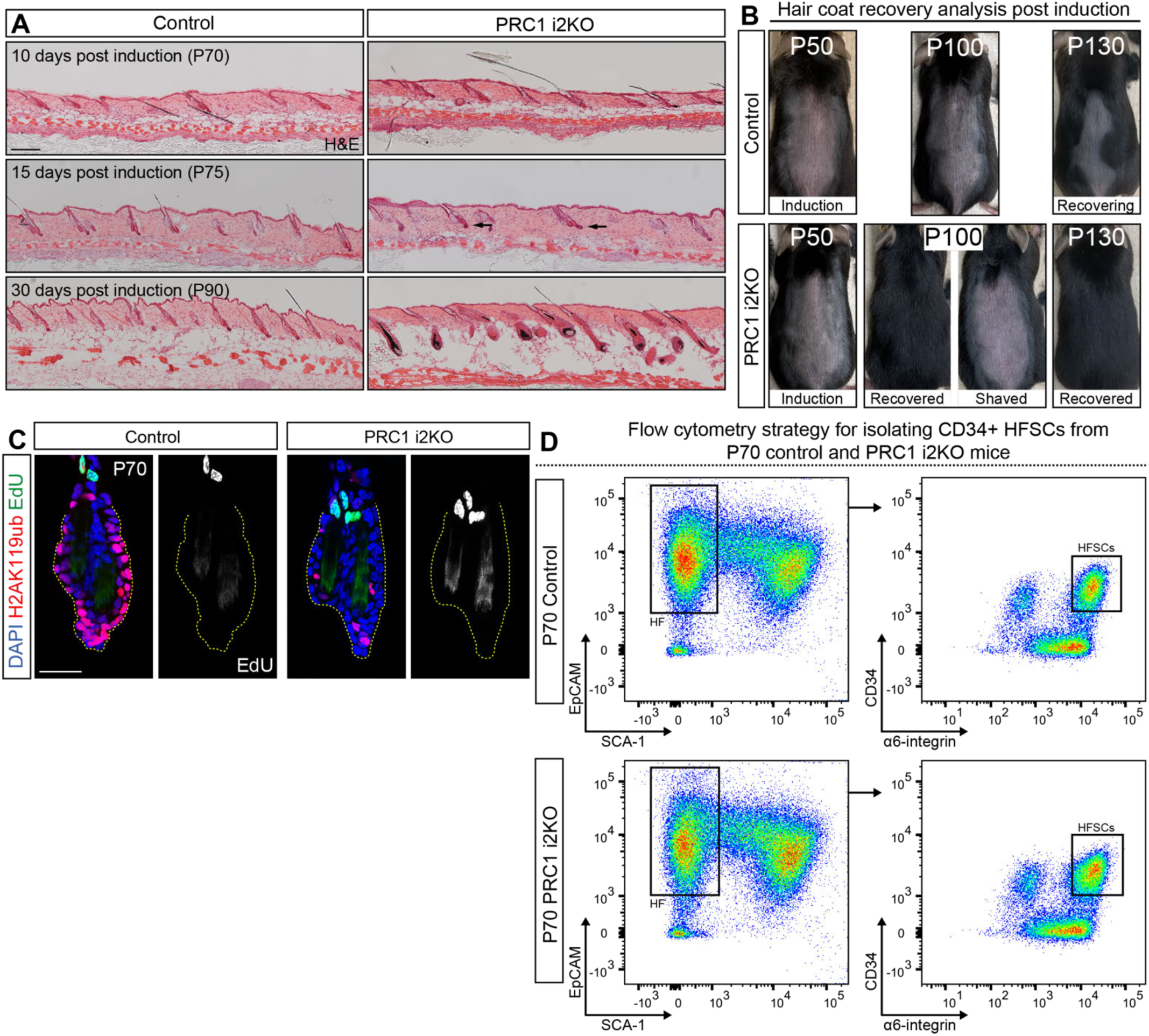
(Related to Fig. 2) Loss of PRC1-mediated H2AK119ub in the bulge induces premature anagen onset. (**A**) H&E analysis of skins from control and PRC1 i2KO back skin 10 days (P70), 15 days (P75), and 30 (P90) days after induction treatment, respectively. (**B**) Images of back skin of control and PRC1 i2KO mice at start of induction (P50), 40 days (P100), and 70 (P130) days after induction treatment, respectively. Hair coat was recovered twice in PRC1 i2KO mice compared to control littermates. (**C**) Epidermal whole-mount IF analysis of H2AK119ub (red), EdU (green), and DAPI (blue) in HF bulges of P70 control and PRC1 i2KO mice. EdU channel shown in grayscale. (**D**) Flow cytometry strategy of isolating HFSCs identified as EpCAM^+^, Sca1^-^, α6-integrin^high^, and CD34^+^ from the back skin of P70 control and PRC1 i2KO mice. Scale bar for IF images: 25μm; for H&E images: 100μm.

**Supplemental Figure 5.**
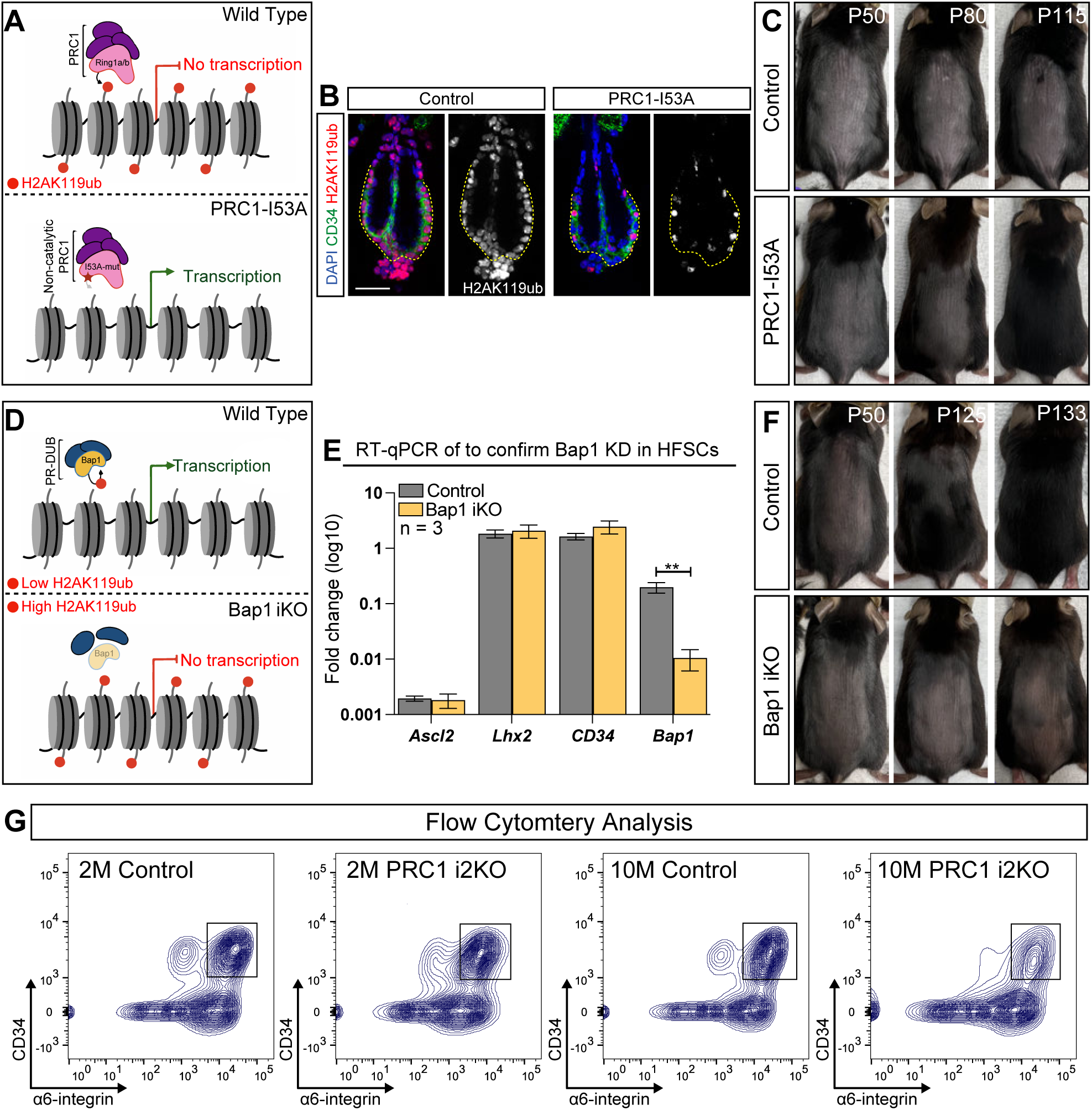
(Related to Figs. 2 & 3) Dysregulation of H2AK119ub levels in the bulge impacts HFSC proliferative potential. (**A**) Schematic illustrating that I53A point-mutation in RING1B ablates its H2AK119ub depositing catalytic function. (**B**) Epidermal whole-mount IF analysis of H2AK119ub (red), CD34 (green), and DAPI (blue) in HFSCs of P70 control and PRC1-I53A HF bulges. H2AK119ub channel shown in grayscale. (**C**) Images of back skin of control and PRC1-I53A mice at start of induction (P50), 20 days (P80), and 55 days (P115) after induction treatment, respectively. (**D**) Schematic illustrating that ablation of BAP1 function in the HFSCs leads to maintenance of H2AK119ub. (**E**) RT-qPCR analysis of epidermal marker (*Ascl2*), HFSC makers (*Lhx2* and *Cd34*) and Bap*1* mRNA in P70 control and Bap1 iKO HFSCs. An unpaired t-test was conducted (n=3 animals from at least two independent litters). P-value for *Bap1* = 0.0015 (**F**) Images of back skin of control and Bap1 iKO mice at start of induction (P50), 65 days (P125), and 73 days (P133) after induction treatment, respectively. (**G**) Contour plots of EpCAM^+^, Sca1^-^, α6-integrin^high^and CD34^+^ HFSCs from the back skin of 2M control and PRC1 i2KO and 10M control and PRC1 i2KO mice. Scale bar: 25μm.

**Supplemental Figure 6.**
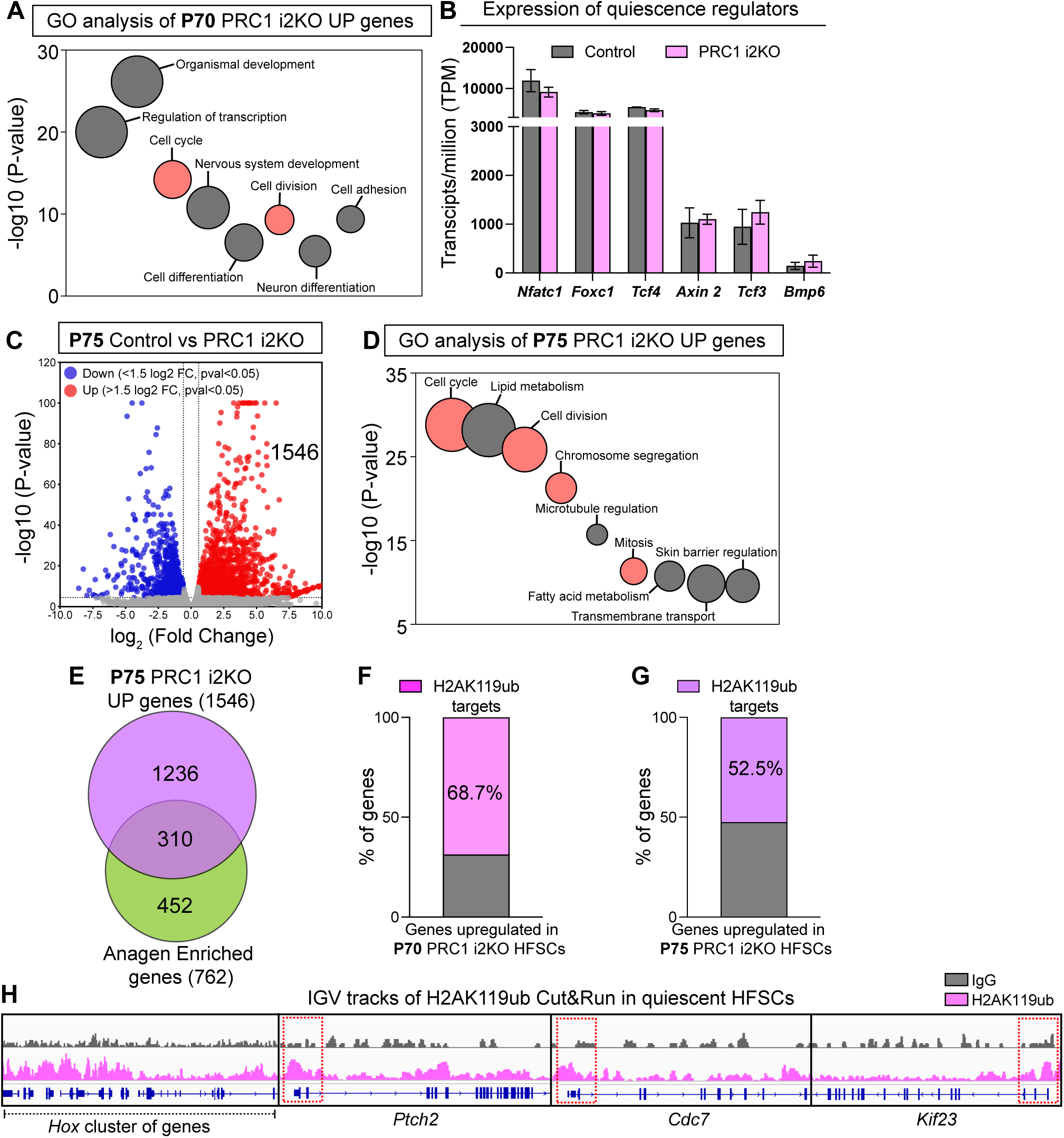
(Related to Fig. 4) Characterization of the transcriptional changes in HFSCs upon H2AK119ub ablation. (**A**) Gene Ontology (GO) analysis of the upregulated genes in P70 PRC1 i2KO HFSCs. (**B**) Graph showing the RNA levels of HFSC quiescence regulators in P70 control and PRC1 i2KO HFSCs. (**C**) Volcano plot showing the differentially expressed genes in FACS-purified HFSCs from P75 control and PRC1 i2KO mice. Genes with absolute fold change ≥ 1.5 and adjusted p-value < 0.05 were considered significantly upregulated or downregulated for further analysis. RNA-seq analysis was done on three biological replicates for each group from at least two independent litters. (**D**) Gene Ontology (GO) analysis of the upregulated genes in P75 PRC1 i2KO HFSCs. (**E**) Venn diagram showing that 310 anagen enriched genes were upregulated in P75 PRC1 i2KO HFSCs. (**F**) Stacked graph showing that 68.7% of upregulated genes in P70 PRC1 i2KO HFSCs are H2AK119ub targets. (**G**) Stacked graph showing that 52.5% of upregulated genes in P75 PRC1 i2KO HFSCs are H2AK119ub targets. (**H**) IGV tracks showing H2AK119ub occupancy on known PRC1 targets (*Hox* cluster) and proliferation promoting and cell cycle genes in quiescent HFSCs.

**Supplemental Figure 7.**
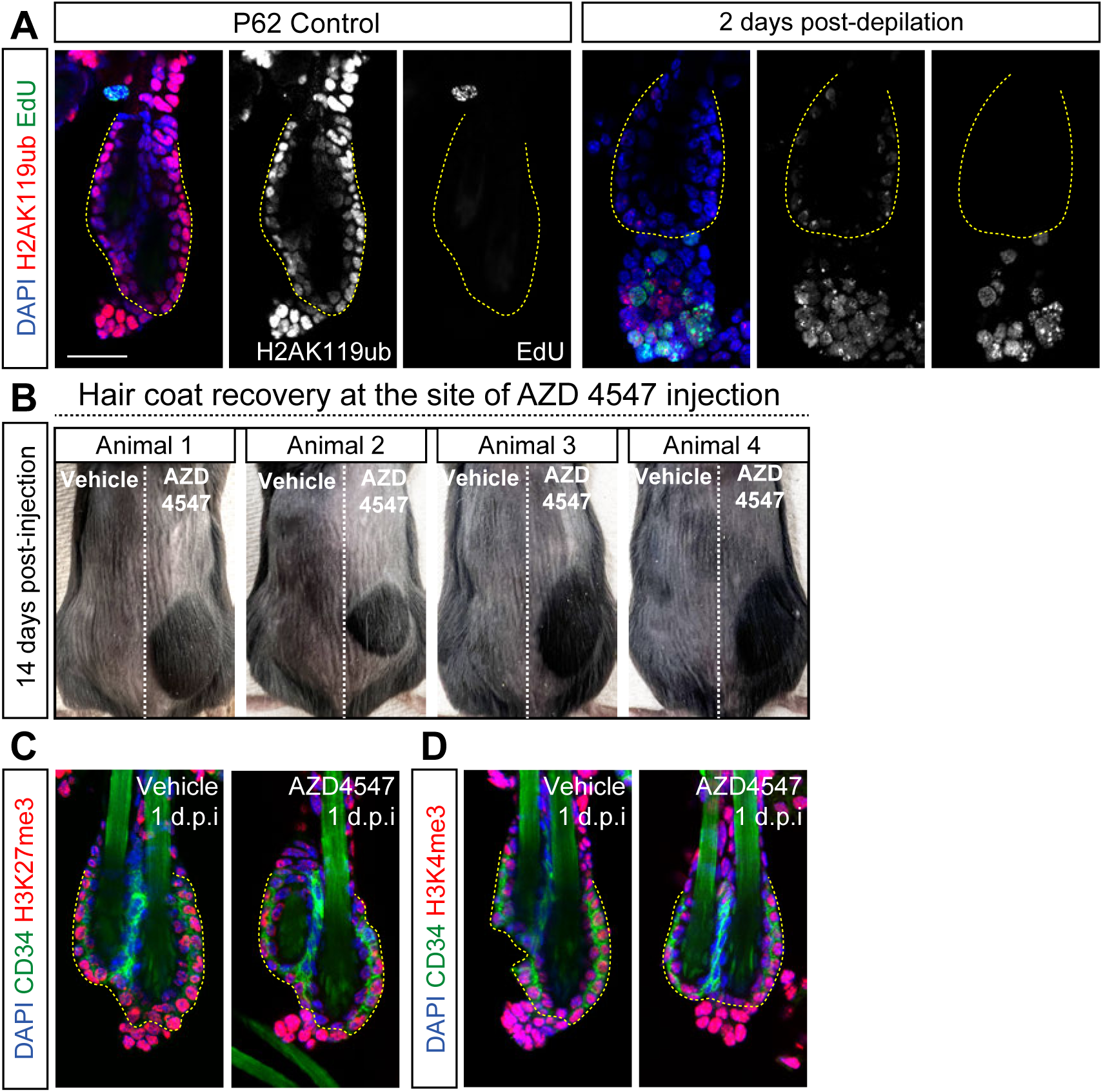
(Related to Fig. 5) Inhibitory FGF signaling maintains H2AK119ub levels in HFSCs. (**A**) Epidermal whole-mount IF analysis of H2AK119ub (red), EdU (green), and DAPI (blue) in skin samples from control (P62) and 2 days post-depilated skin showing that EdU+ proliferating bulges of depilated skin have markedly lower levels of H2AK119ub when compared to control skin. H2AK119ub and EdU channels shown in grayscale. (**B**) Representative images of back skin of animals 14 days after intradermally injected with either vehicle or FGFR inhibitor, AZD4547. (**C**) Epidermal whole-mount IF analysis of H3K27me3 (red), CD34 (green), and DAPI (blue) showing no changes in H3K27me3 levels in HFSCs of vehicle or AZD4547 injected site. (**D**) Epidermal whole-mount IF analysis of H3K4me3 (red), CD34 (green), and DAPI (blue) showing no changes in H3K4me3 levels in HFSCs of vehicle or AZD4547 injected site. Scale bar: 25μm.

## Materials and Methods

### Mouse strains

Research within this publication complies with relevant ethical regulations. All experimental protocols using animals was approved by and in accordance with the Institutional Animal Care and Use Committee (IACUC) guidelines (Protocol No. LA11-00020) at the Icahn School of Medicine at Mount Sinai (ISSMS).

Mice were housed in the Center for Comparative Medicine and Surgery at Icahn School of Medicine at Mount Sinai (ISMMS) in accordance with the Institutional Animal Care and Use Committee. Mice were genotyped by PCR using ear skin DNA that was extracted using DirectPCR Lysis Reagent (Viagen Biotech Inc), according to manufacturer’s instructions. *K15-CrePR* (005249) and *Bap1^flox/flox^* (031874) ^mice^ lines were obtained from The Jackson Laboratory. We were gifted the *Ring1a*-null and *Ring1b ^flox/flox^* mice by Dr. Bickmore and *Ring1b^I53A^*mice by Dr. Vidal, respectively. C57BL6/J (wild type) mice were acquired from either The Jackson Laboratory or Charles River Laboratories. Both male and female mice were used in this study.

### Chemical induction and intradermal injections

For RU-486 (Cayman Chemicals) topical treatment to induce PRC1 ablation, I53A-mutation, and Bap1 ablation, RU-486 was dissolved in 100% ethanol (Sigma-Aldrich) to a final concentration of 10 mg/ml. 100μl of RU-486 was topically applied on shaved dorsal skin of postnatal day 50 (P50) once per day for 10 days. Control mice were treated with the same amount of 100% ethanol. For intradermal injections, 5mg of AZD4547 (Selleckchem, S2801) was dissolved in 1mL DMSO and 4mL of sterile PBS to reach a final concentration of 1mg/mL. 100μl of this solution was injected intradermally in the shaved dorsal skin of C57BL6/J (wild type) for 5 consecutive days. Control side was injected with 100μl of Vehicle (PBS+ 1% DMSO) solution.

### Hair cycle analysis and depilation studies

Hair cycle progression was documented by standardized photographs at the start of each experiment and bi-weekly thereafter. For testing hair coat recovery, the back skin of mice was shaved with an electric clipper. Anagen onset was determined by darkening of the skin followed by hair growth as previously described^72^. Once the hair coat recovery reached ∼90% of the back skin, the mice were shaved again to monitor the entry into next anagen. Days taken to recover hair coat was calculated after the induction treatment was completed at P60 and was observed in three- four biological replicates from at least two independent litters. Graphs were generated using GraphPad Prism software (v.9.5.1). An unpaired t-test was conducted, and 0.05 level of confidence was accepted for statistical significance. ** indicates p-values < 0.001.

For depilation studies, back skin of telogen C57BL6/J (wild type) mice was shaved and depilated using wax strips (Sally Hensen). Skin samples were collected 48 hours after depilation for epidermal whole mount IF assay.

### Immunofluorescence staining and microscopy

For epidermal whole mount immunofluorescence (IF) we followed a previously described protocol^73^ with a few variations as follows. Skin samples were collected and the adipose layer from the dorsal side was scraped off using a surgical scalpel. The scraped skin sample was washed with phosphate-buffered saline (PBS) before cut into 4mm x 4mm square sections and incubated for 5 hours in 1.5 mL Eppendorf tubes with 1mg/mL Dispase (Gibco, 17-105-041) and 20mM EDTA (Corning, 46-034-CI) solution in Dulbecco’s Phosphate-Buffered Saline (DPBS) (Fisher Scientific, MT21031CV) at 37°C on a shaker set at 100 rotations per minute (rpm). After incubation, the epidermal layer was carefully peeled away from the dermis using a S-shaped tweezer and were fixed in 4% paraformaldehyde (PFA; Electron Microscopy Sciences) in PBS on a nutator. After fixing, tissue samples were washed twice with PBS for 10 minutes (min). Samples could be stored for up to three weeks at 4°C. If proceeding with the assay samples were permeabilized with 0.5% Triton-X in PBS for 30 min on a nutator. Following permeabilization, samples were washed twice with PBS for 10 min and blocked for 2 hours at RT with blocking solution [1% Triton X- 100, 1% bovine serum albumin (BSA), 0.01% Gelatin and 0.25% normal donkey serum]. Primary antibodies (**Table 1**, below) were diluted in blocking solution, and incubations were carried out overnight at 4°C, followed by incubation in secondary antibodies (**Table 1**, below) for 2 hours at RT. Slides were counterstained with 4′,6-diamidino-2-phenylindole (DAPI) (Sigma, 32670-5MG-F) and mounted using antifade mounting media.

**Table 1.** List of primary and secondary antibodies used in this study.

| Antibody | Vendor | Catalog Number |
| --- | --- | --- |
| Rabbit anti-H2AK119ub | Cell Signaling | 8240S |
| Rabbit anti-H3K4me3 | Abcam | ab8580 |
| Rabbit anti-H3K27me3 | Cell Signaling | 9733S |
| Rat anti-CD34 | eBiosciences | 14-0341-82 |
| Rabbit anti-SOX9 | Abcam | ab185966 |
| Rabbit anti-Activated CASAPASE 3 | R&D systems | AF835 |
| Rabbit anti-P19<sup>ARF</sup> | Abcam | ab80 |
| Anti-rat Alexa Fluor 488 | Jackson ImmunoResearch Labs | 712-545-150 |
| Anti-rat Alexa Fluor 647 | Jackson ImmunoResearch Labs | 712-605-150 |
| Anti-rabbit Alexa Fluor 488 | Jackson ImmunoResearch Labs | 711-545-152 |
| Anti-rabbit Alexa Fluor 594 | Jackson ImmunoResearch Labs | 711-585-152 |
| Anti-rabbit Alexa Fluor 647 | Jackson ImmunoResearch Labs | 711-605-152 |

For IF staining of skin sections, skin samples were collected and embedded into Tissue-Tek optimal cutting temperature (OCT) compound blocks (Electron Microscopy Sciences, 62550-12) and cut into 9-μm sections using a Leica Cryostat (CM3050-S). Slides were either stored at 80°C or used to proceed with the assay. Slides were fixed for 10 min in 4% PFA in PBS. Following couple of washes with PBS, the sections were outlined with a hydrophobic barrier pen (ImmEdge PAP Pen, Vector Laboratories, H-4000) and blocked for 2 hours at RT or overnight at 4°C in PBS with blocking solution. Primary antibodies (**Table 1**, below) were diluted in blocking solution, and incubations were carried out for 1 hour at RT or overnight at 4°C, followed by incubation in secondary antibodies (**Table 1**, below) for 2 hours at RT. Slides were counterstained with DAPI and mounted using antifade mounting media.

Slides were imaged using Leica DM5500 slide microscope (Leica Microsystems GmbH, Wetzlar, Germany) with 40X objective. Epidermal wholemount samples were imaged using the LAS X software (Leica Application Suite X 4.3.0.24308) on Leica Stellaris 8 (Leica Microsystems GmbH, Wetzlar, Germany) confocal microscope with the 40x/1.3 HC PlanApo oil immersion lens (Leica Microsystems GmbH, Wetzlar, Germany). Images were processed using the NIH ImageJ software (v2.14.0/1.54f) and are presented as maximum intensity projection images or a single Z stack. Images were further processed and assembled into panels using Adobe Photoshop (v.20.0.3) and Adobe Illustrator (v.24.0).

### EdU pulse-chase experiment

Mice were injected with 125 μg of EdU (Thermo Fisher, E10187) in solution. Skin samples were collected 2 hours after injection and were either embedded in OCT or collected for epidermal whole mount IF. Slides or whole mount samples were fixed and permeabilized as described above, followed by a 30-min incubation with a Click-iT reaction cocktail (Thermo Fisher Scientific, C10340 or C10637). Samples were washed twice with 3% BSA before proceeding with IF assay as described above. HFs positive for EdU incorporation in the bulge and hair germ region was considered activated and percentage of proliferative HFs was determined by counting 50 HFs in each biological replicate across all time points. Graphs were generated using GraphPad Prism software (v.9.5.1). An unpaired t-test was conducted, and 0.05 level of confidence was accepted for statistical significance. ** indicates p-values < 0.001 and **** indicates p- values < 0.00001. Primer details are in **Table 2**, below.

**Table 2.**
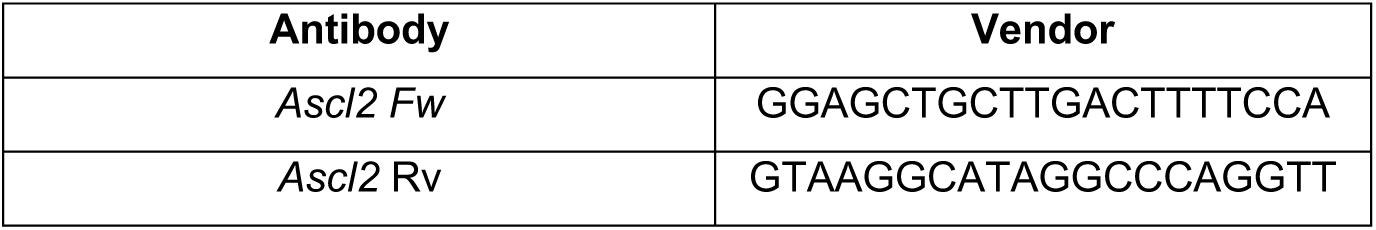
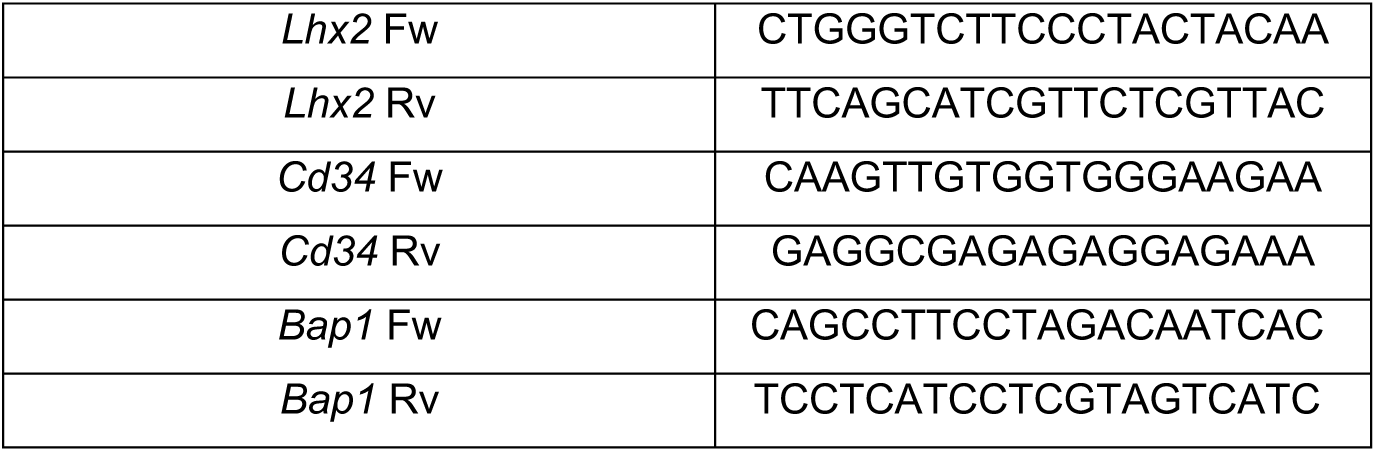
List of primers for qPCR (Fw: forward; Rv: reverse).

### Nuclear Flow Cytometry

For detection of histone modifications in HFSCs of mice of different ages, P21, P23, P26, and P55, all time points were analyzed at the same time, and the experiment was repeated with three independent litters. The back skin from adult mice was collected and the adipose layer from the dorsal side was scraped off using a surgical scalpel. The scraped skin sample was washed with PBS prior to incubation with 0.25% Trypsin/EDTA (Fisher, 25-053-CI) at 37°C for 1 hour on a rotation plate set at 60 rpm. After incubation, the epidermal cells, including the HFSCs, were scraped off from the trypsinized skin into the plate. 25mLs of E-media was added to the cell suspension and was then sequentially strained through 100μm and 40μm filters. The cell pellet was washed twice with DPBS. Cells were stained with Zombie Violet™ Fixable Viability Kit (Biolegend, 423114) for 10 min on ice. Cells were washed twice with DPBS prior to fixing using fresh solution with a final concentration of 1% formaldehyde (Thermo Fisher Scientific, 28906) for 10 min at RT. Crosslinking was stopped by the addition of Glycine (final concentration 125mM). Cells were washed twice with DPBS before freezing in 90%FBS/10%DMSO. Once HFSCs from all experimental samples were collected, the frozen cell suspension was thawed on ice and washed with DPBS. Cells were first stained with 1:200 PE-Sca1 (Biolegend, 108108), 1:200 APC/Cy7-α6-integrin (Biolegend, 313628) and 1:20 FITC- CD34 (Thermo Fisher, 11-0341-82) in staining buffer (HBSS + 2% Fetal Bovine Serum) for 30 minutes on ice and then washed twice with DPBS. Cells were then permeabilized with 200μl of permeabilization buffer (0.2% Triton X-100 in DPBS) for 15 min at RT. Primary antibodies (histone modifications listed in Table 1) were added directly at 1:1000 dilution into the permeabilizing cells and further incubated for 25 minutes at RT. Cells were washed once with 2mL of permeabilization buffer, before resuspending in 200μl of permeabilization buffer with secondary antibodies (Anti-rabbit Alexa Fluor 647 conjugate) diluted at 1:500 concentration and incubated for another 25 minutes at RT. A fraction of the cells were also stained with just secondary (2°) antibodies to serve as negative control. Cells were washed twice with staining buffer before proceeding with analysis using a benchtop flow cytometer (BD) in the Flow Cytometry Core Facility at ISMMS. HFSCs were identified as Sca1(-), α6-integrin (high), and CD34(+). FACS strategy plots were generated using FlowJo portal (10.10.0), and percentage of of histone modification (+) HFSCs were identified. Graphs were generated using GraphPad Prism software (v.9.5.1). Ordinary one-way ANOVA (Dunnett’s multiple comparisons test) was conducted, and 0.05 level of confidence was accepted for statistical significance. * indicates p-values < 0.01 and ** indicates p-values <0.001.

### Fluorescence-activated cell sorting (FACS)

FACS isolation of HFSCs from the back skin of control, PRC1 i2KO, PRC1-I53A and Bap1 iKO mice was performed as follows. The back skin from adult mice was collected and the adipose layer from the dorsal side was scraped off using a surgical scalpel. The scraped skin sample was washed with PBS prior to incubation with 0.25% Trypsin/EDTA at 37°C for 1 hour on a rotation plate set at 60 r.p.m. After incubation, the epidermal cells, including the HFSCs, were scraped off from the trypsinized skin into the plate. 25mLs of E-media was added to the cell suspension and was then sequentially strained through 100μm and 40μm filters. The cell pellet was washed twice with DPBS before proceeding with staining with cell surface markers. The cells were stained with 1:200 PerCP-Cy5.5-Sca1 (Thermo Fisher Scientific, 45-5981-82), 1:200 FITC-α6-integrin (eBioscience, 11-0493-81), 1:100 APC-EpCAM (Biolegend, 324207) and 1:20 Alexa700-CD34 (eBioscience, 56-0341-82) in staining buffer (HBSS + 2% Fetal Bovine Serum) for 30 minutes on ice and then washed twice with DPBS before cell sorting. HFSCs were sorted by gating on EpCAM (+), Sca1(-), α6-integrin(high) and CD34(+). All cell isolations were performed on a FACS Influx instrument (BD) in the Flow Cytometry Core Facility at ISMMS. FACS strategy plots were generated using FlowJo portal (10.10.0). Additionally, percentage of HFSCs at different ages in control and PRC1 i2KO mice was determined and graphs were generated using GraphPad Prism software (v.9.5.1). Ordinary one-way ANOVA (Dunnett’s multiple comparisons test) was conducted, and 0.05 level of confidence was accepted for statistical significance. ** indicates p-values <0.001.

### Colony formation assay

FACS-purified HFSCs were plated on mitomycin C (Fisher Scientific, BP25312)-treated J2 fibroblast feeders at a density of 10,000 cells/well in12-well plates in E media supplemented with 15% (v/v) serum and 0.3 mM calcium as described in previous studies^74^. After significant colonies were formed either in control or KO wells, cells were fixed and stained with 1% (wt/vol) Rhodamine B (Millipore Sigma, R6626). Colony diameter was measured from scanned images of plates using ImageJ (v2.14.0/1.54f) and percentage area covered by colonies was calculated. Graphs were generated using GraphPad Prism software (v.9.5.1) and unpaired t-test was conducted, and 0.05 level of confidence was accepted for statistical significance. ** indicates p-values < 0.001 and *** indicates p-values <0.0001.

### RNA purification, RT-qPCR, and RNA-sequencing library preparation

A total of 50,000 FACS-purified cells were collected directly into the RLT Plus buffer (QIAGEN, 1053393), and RNA was purified from these sorted cells with the RNeasy Plus Micro Kit (QIAGEN, 74034) and DNase1 treatment (QIAGEN, 79254) according to the manufacturer’s instructions. cDNA was reverse transcribed from total NA using qScript cDNA Supermix (Quanta Biosciences, 101414-106) according to manufacturer’s instructions. Samples were analyzed by RT-qPCR using LightCycler 480 SYBR Green I Master Mix (Roche) on a Roche Lightcycler 480 instrument.

For RNA-seq library preparation, sample quality was measured using an Agilent Bioanalyzer. Only samples with RNA integrity numbers of >8 were used for library preparation. For generating RNA-seq libraries, 50 ng of total RNA was subjected to polyadenylate selection using Universal Plus mRNA-Seq with NuQuant (TECAN, 0520-A01). Following elution, mRNA was subjected to fragmentation at 94°C for 8 min. First strand, second-strand cDNA synthesis, end repair, adapter ligation, and amplification were carried out by following the manufacturer’s instructions.

### RT-qPCR analysis and strategy

Fold change in mRNA levels of *Ascl2, Lhx2, Cd34,* and *Bap1* in HFSCs isolated from P70 control and Bap1 iKO mice was calculated by first averaging the Ct values of technical replicates from each trial. ΔCt was calculated by subtracting *Gapdh* Ct average from the Ct average of AEGs. Then the of the 2^- ΔCt was calculated for each trial. Fold change was then calculated by dividing Bap1 iKO 2^- ΔCt by control 2^- ΔCt. Bar graph is presented as mean ± SD. Three animals for each group from at least two independent litters were used. Graphs were generated using GraphPad Prism software (v.9.5.1). An unpaired t-test was conducted, and 0.05 level of confidence was accepted for statistical significance. ** indicates p-values < 0.001. Primer details are in **Table 2**, below.

### RNA-seq analysis and data visualization

Paired-end RNA-seq reads were aligned to the mouse reference genome (mm39) using STAR aligner (v2.7.9a)^75^. Three samples each for control and PRC1 i2KO were processed but one control in P70 and one in P75 samples were excluded for further analysis due to low quality. Gene expression levels (from the Ensembl annotation v105, both coding and noncoding) were quantified using the RSEM software (v1.3.3) to estimate counts and TPMs (transcript per million)^76^. Differential expression analysis was performed for protein-coding genes using DESeq2 (v1.38.2)^77^ after excluding genes with TPM < 1 in all samples. Genes reaching *P* value < 0.05 and log2-transformed fold change > 1.5 (or < -1.5) were considered to be significantly differentially expressed. Volcano plots were generated using the SRPlot web tool^78^.

### Gene ontology and Gene Set Enrichment Analysis

Gene Ontology (GO) enrichment analysis was performed using the database for annotation, visualization and integrated discovery (DAVID) web-accessible tool^79^. For GSEA analysis of PRC1 i2KO HFSCs, all protein-coding genes were ranked by multiplication of their fold change direction with -log10(p-value) and this pre-ranked gene list was used as input for the GSEA tool (GSEA_4.1.0). The gene set for GSEA was a previously defined Anagen enriched gene list^42^.

### Cleavage Under Targets & Release Using Nuclease (Cut&Run) analysis and data visualization

120,000 FACS-enriched quiescent HFSCs were isolated from the back skin of P55 C57BL/6 wildtype mice. Cut&Run was conducted using the CUTANA™ ChIC/CUT&RUN Kit (Epicypher, 14-1048) following a previously described protocol optimized for FACS-isolated epidermal keratinocytes^80^. Sequencing libraries were generated using NEBNext Ultra II DNA Library Prep Kit for Illumina (NEB, E7645L) and NEBNext Multiplex Oligos (NEB, E6609L) for Illumina following the manufacturer’s instructions. PCR-amplified libraries were purified using 1 X ratio of Agencourt AMPure-XP Beads (Beckman) and eluted in 15μl TE buffer.

All libraries were sequenced on an Illumina NextSeq using 150 bp paired-end reads. Reads were trimmed with Trim Galore (v0.6.7) and aligned to the mouse reference genome (mm10) using Bowtie2 (v.2.2.5). Deduplicated reads were removed using samtools (v.1.14). Peaks were identified using a sliding window method to determine genomic regions with >1.5 mapping read enrichment over IgG control and adjusted p < 0.05. The BAM file for each replicate was converted to a TDF file for Integrative Genomics Viewer (IGV) software.

## Acknowledgements

We thank Sergei Ezhkov and all other Ezhkova lab members for their help and critical suggestions. We would like to thank Dr. Miguel Vidal for the *Ring1a*-null and *Ring1b-floxed* mice and Dr. Wendy A. Bickmore for the *Ring1b^I53A^*mice. We would also like to thank the Flow Cytometry Core and the Microscopy Core facilities at Icahn school of Medicine at Mount Sinai. Some graphical illustrations in this manuscript were generated using Biorender.com.

## Contributions

Conceptualization, P.F. and E.E.; methodology, P.F., D.Z., and E.E.; investigation, P.F., M.Y.L., Y.Z., M.M., X.Z., and P.M.G.; resources, D.Z., and E.E.; writing – original draft, P.F. and E.E.; writing – review and editing, P.F., M.Y.L., Y.Z., M.M., X.Z., D.Z, and E.E.; project administration, P.F. and E.E.; funding acquisition, P.F. and E.E.

## Data Availability

All data needed to evaluate the conclusions in the paper are present in the paper and/or the Supplementary Materials. The accession number for the bulk RNA-seq comparing Telogen and Anagen HFSCs to identify anagen enriched genes is GSE172082. The accession number for the bulk RNA-seq and CUT&RUN data generated in this study will be submitted to Gene Expression Omnibus (GEO) upon publication. All other data are available from the corresponding author on reasonable request.

